# Effect of Charge Distribution on the Dynamics of Polyampholytic Disordered Proteins

**DOI:** 10.1101/2022.07.04.498718

**Authors:** Dinesh Sundaravadivelu Devarajan, Shiv Rekhi, Arash Nikoubashman, Young C. Kim, Michael P. Howard, Jeetain Mittal

## Abstract

The stability and physiological function of many biomolecular coacervates depend on the structure and dynamics of intrinsically disordered proteins (IDPs) that typically contain a significant fraction of charged residues. Although the effect of relative arrangement of charged residues on IDP conformation is a well-studied problem, the associated changes in dynamics are far less understood. In this work, we systematically interrogate the effects of charge distribution on the chain-level and segmental dynamics of polyampholytic IDPs in dilute solutions. We study a coarse-grained model polyampholyte consisting of an equal fraction of two oppositely charged residues (glutamic acid and lysine) that undergoes a transition from an ideal chain-like conformation for uniformly charge-patterned sequences to a semi-compact conformation for highly charge-segregated sequences. Changes in the chain-level dynamics with increasing charge segregation correlate with changes in conformation. The chain-level and segmental dynamics conform to simple homopolymer models for uniformly charge-patterned sequences but deviate with increasing charge segregation, both in the presence and absence of hydrodynamic interactions. We discuss the significance of these findings, obtained for a model polyampholyte, in the context of a charge-rich intrinsically disordered region of the naturally occurring protein LAF-1. Our findings have important implications for understanding the effects of charge patterning on the dynamics of polyampholytic IDPs in dilute conditions using polymer scaling theories.

## 1 Introduction

Intrinsically disordered proteins (IDPs) or regions (IDRs) are a driving force in the formation of membraneless organelles^1,2^ that are responsible for many physiological functions, *e*.*g*., signaling, stress response, transcription, and metabolic processes.^3–7^ Due to the lack of well-defined three-dimensional folded structures, IDPs display remarkable heterogeneity in terms of structural architectures, binding multiplicity, and functional diversity. Many known IDPs are polyampholytes, ^8^ a class of polymers that consists of both positively and negatively charged monomers in varying fractions. Naturally occurring polyampholytes include gelatin and bovine serum albumin, but synthetic polyampholytes have also been made. ^9^ Many theoretical and computational studies have investigated the conformations and phase behavior of polyampholytes.^10–18^ Among these studies, Samanta *et al*.^17^ showed that randomly patterned polyampholytes exhibit a non-monotonic trend in their radius of gyration *R*_g_ with an increase in the inverse Debye screening length. Using molecular dynamics (MD) simulations and scaling theory, Wang and Rubinstein^16^ identified three well-defined regimes in the coil–globule transition of a diblock polyampholyte with increasing electrostatic interaction strength: in the first regime, the chain folds through the overlap of the two oppositely charged blocks; in the second regime, the chain collapses to a globule with densely packed charged sections; and in the third regime, oppositely charged monomers tightly bind in a cascade of multipole formation.

The conformations of polyampholytic IDPs depend on several parameters related to electrostatic interactions such as the fraction of charged residues, ^19,20^ the net charge per residue,^21^ and the specific arrangement of charges in a sequence. ^19^ For example, charge patchiness has recently been shown to dictate the conformations adopted by polyampholytes. ^18^ To reduce this complex parameter space, it is desirable to find composite metrics that can be used to characterize and predict polyampholyte properties.^19,22,23^ Sawle and Ghosh^22,23^ proposed the sequence charge decoration (SCD) parameter that successfully described the trends in conformations adopted by single-chain polyampholytic IDPs with increasing charge segregation. In addition, the critical temperature of polyampholytic condensed phases has been shown to correlate linearly with SCD. ^24^ These and other studies also revealed that single-chain properties are indicative of the propensity of IDP solutions to phase separate. ^24,25^

Despite extensive literature on the structural properties of polyampholytes, studies on their dynamics and how they respond to changes in charge distribution are scarce. A detailed study on single-chain polyampholyte dynamics is an important first step towards understanding the dynamics and rheology of IDPs in dilute and condensed phases and, ultimately, facilitating the design of tailored protein sequences with specific dynamical responses and material properties. For example, the translational dynamics of polyampholytes in the condensed phase was recently shown to depend on the degree of charge segregation in the sequence.^26^ In this paper, we perform molecular simulations to investigate the effect of charge distribution on the single-chain dynamics of a model polyampholyte. We use a coarse-grained (single bead per residue) model IDP consisting of an equal fraction of negatively charged glutamic acid (E) and positively charged lysine (K) residues. Previously, coarse-grained models have been successfully used to investigate the molecular forces that govern the phase behavior and assembly of IDPs. For example, coarse-grained models were instrumental in revealing the effect of charge patterning in LAF-1 RGG phase separation by itself^27^ and in the presence of polynucleotides,^28^ as well as in establishing the relationship between sequence specificity and conformational properties of IDPs. ^29^ Our results reveal that the conformational and dynamical properties associated with the end-to-end vector of polyampholytes show a similar dependence on charge patterning. However, we find that the applicability of homopolymer models for describing the correlation between structure and dynamics in polyampholytes depends on the extent of charge segregation within them. Specifically, the dynamics of uniformly charge-patterned sequences are well-described by the Rouse (Zimm) model in the absence (presence) of hydrodynamic interactions, but the dynamics of highly charge-segregated sequences deviate from these classic homopolymer models. Finally, we show that our findings also apply to the charge-rich intrinsically disordered LAF-1 RGG domain, which represents a large class of naturally occurring sequences with uniform charge distribution.^19^

The rest of this paper is organized as follows. The molecular model and simulation methodology are presented in Section 2. The conformational and dynamical properties of the model polyampholyte are presented and analyzed in Sections 3.1 and 3.2, respectively. These findings are then discussed in the context of a charge-rich IDR of the naturally occurring protein LAF-1 RGG in Section 3.3. The effects of hydrodynamic interactions on the dynamics of both the model polyampholyte and LAF-1 RGG are discussed in Section 3.4 before the results are summarized in Section 4.

## 2 Molecular Model and Simulation Details

Our model polyampholyte was a linear chain of E and K residues represented as single beads and bonded to form a specified sequence of *N* total residues. We primarily investigated chains with *N* = 50 residues, but we also performed simulations with *N* = 30 and *N* = 70 residues. All chains consisted of an equal number of E and K residues such that the whole chain had zero net charge. The degree of charge segregation in a given sequence was quantified using the SCD parameter,^22^

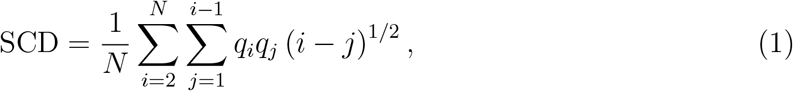

where *q*_*i*_ and *q*_*j*_ are the charges of residues *i* and *j*, respectively, in a given sequence. The SCD is maximum for a perfectly alternating sequence (−0.413 for *N* = 50) and minimum for a diblock sequence (−27.842 for *N* = 50). To facilitate comparison with other polyampholyte compositions and lengths, we will use a normalized SCD (nSCD) parameter that is scaled by these bounds so that nSCD = 0 for a perfectly alternating sequence and nSCD = 1 for a diblock sequence. ^30^ We generated 42 E–K sequences (Figure S1), out of which 15 sequences that spanned the entire range of nSCD were selected for detailed analysis. This subset of E–K variants (EKVs) of length *N* = 50 is shown in Figure 1 with their identifying number and nSCD value.

**Figure 1:**
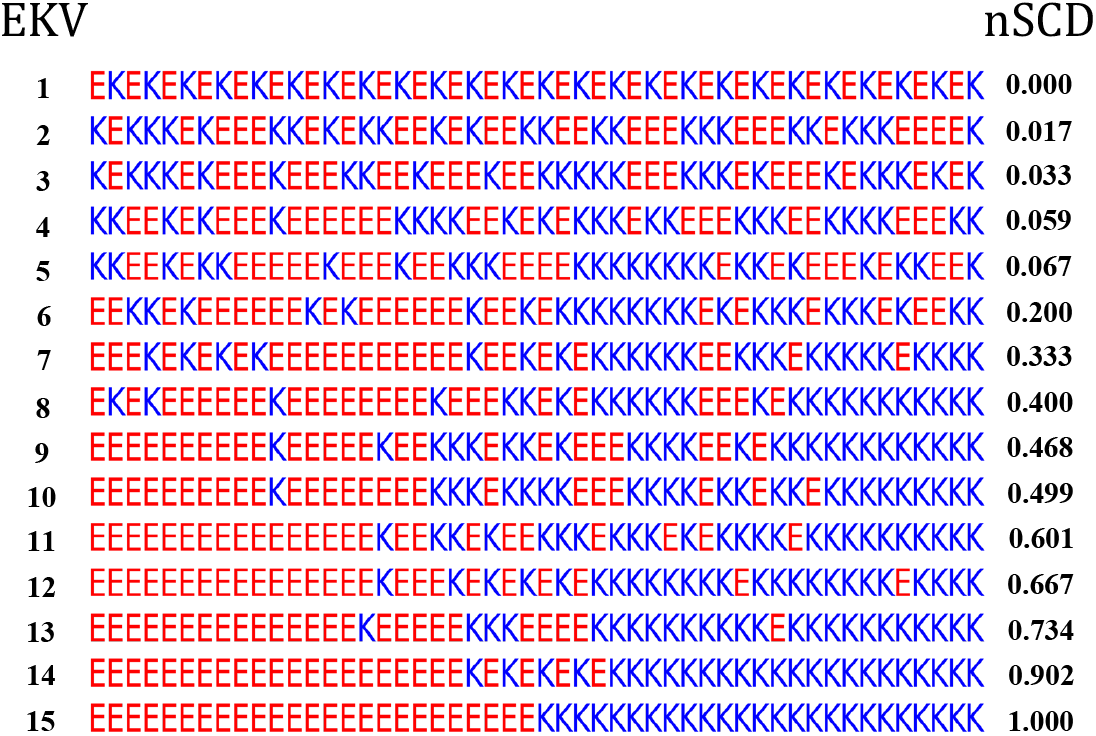
Fifteen selected E–K variants (EKVs) for chain length *N* = 50 with their identifying number and normalized SCD.

Interactions between bonded residues were modeled using a harmonic potential,

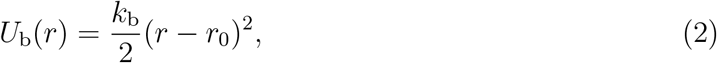

where *r* is the distance between two bonded residues, *k*_b_ = 20 kcal*/* (mol Å^2^) is the spring constant, and *r*_0_ = 3.8 Å is the equilibrium bond length. ^31^ Van der Waals interactions between nonbonded residues were modeled using a modified Lennard-Jones (LJ) potential that allows the attraction between two residues *i* and *j* to be scaled independently from the short-range repulsion, ^32–34^

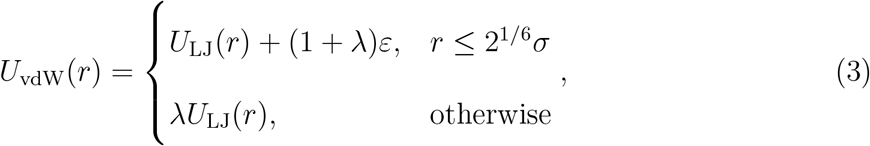

where *U*_LJ_ is the standard LJ potential,

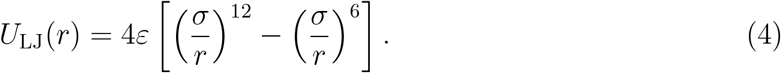

The parameters of *U*_vdW_ are the average hydropathy *λ* = (*λ*_*i*_ + *λ*_*j*_)*/*2 of residues *i* and *j*, the average diameter *σ* = (*σ*_*i*_ + *σ*_*j*_)*/*2 of residues *i* and *j*, and the interaction strength *ε*. The Kapcha–Rossky hydropathy values of 0.460 and 0.514 were used for the E and K residues, respectively.^35,36^ Furthermore, the diameters of the E and K residues were set to 5.92 Å and 6.36 Å, respectively. ^35^ We set the interaction strength to *ε* = 0.2 kcal*/*mol, which was previously found to reproduce the experimentally measured *R*_g_ values of selected IDPs.^35^ The pair potential *U*_vdW_ and its forces were truncated to zero at a distance of 4 *σ* for computational efficiency. The electrostatic interactions between nonbonded residues were modeled using a Coulombic potential with Debye-Hückel electrostatic screening, ^37^

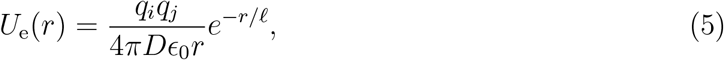

where *D* is the dielectric constant of the medium, *ϵ*_0_ is the permittivity of vacuum, and *ℓ* is the Debye screening length. In our simulations, we used *D* = 80 and *ℓ* = 10 Å, which corresponds to an aqueous solution with physiological salt concentration of approximately 100 mM. The electrostatic potential and its forces were truncated to zero at a distance of 35 Å.

A single chain was placed in a cubic simulation box of edge length 160 Å with periodic boundary conditions applied in all three Cartesian directions. This box size is sufficiently large to prevent unphysical self-interactions between the chain and its periodic images. We performed Langevin dynamics (LD) simulations at temperature *T* = 300 K. We set the residue friction coefficient to *γ*_*i*_ = *m*_*i*_*/t*_damp_, where *m*_*i*_ is the residue mass (129.1 g*/*mol for E and 128.2 g*/*mol for K) and *t*_damp_ = 1000 fs is the damping parameter. These *γ*_*i*_ values were chosen to create weak coupling to a thermostat and were not intended to represent the hydrodynamic forces on the residues from a solvent such as water. All the simulations were carried out for a total duration of 1 *µ*s with a timestep of 10 fs using HOOMD-blue (version 2.9.3) ^38^ with features extended using azplugins (version 0.10.1). ^39^ The simulation trajectories were saved every 1000 fs for characterizing the conformations and chain-level dynamics, and every 100 fs to 1000 fs for characterizing the segmental dynamics. All the measured conformational and dynamical properties were averaged over two independent replicas, with each replica divided into 4 independent blocks to estimate error bars.

## 3 Results and Discussion

### 3.1 Conformations of model polyampholytes

We first characterized the chain size of the EKVs by computing their radius of gyration *R*_g_ = ⟨tr **G**⟩^1*/*2^ (Figure 2a) from the average of the trace of the gyration tensor,

**Figure 2:**
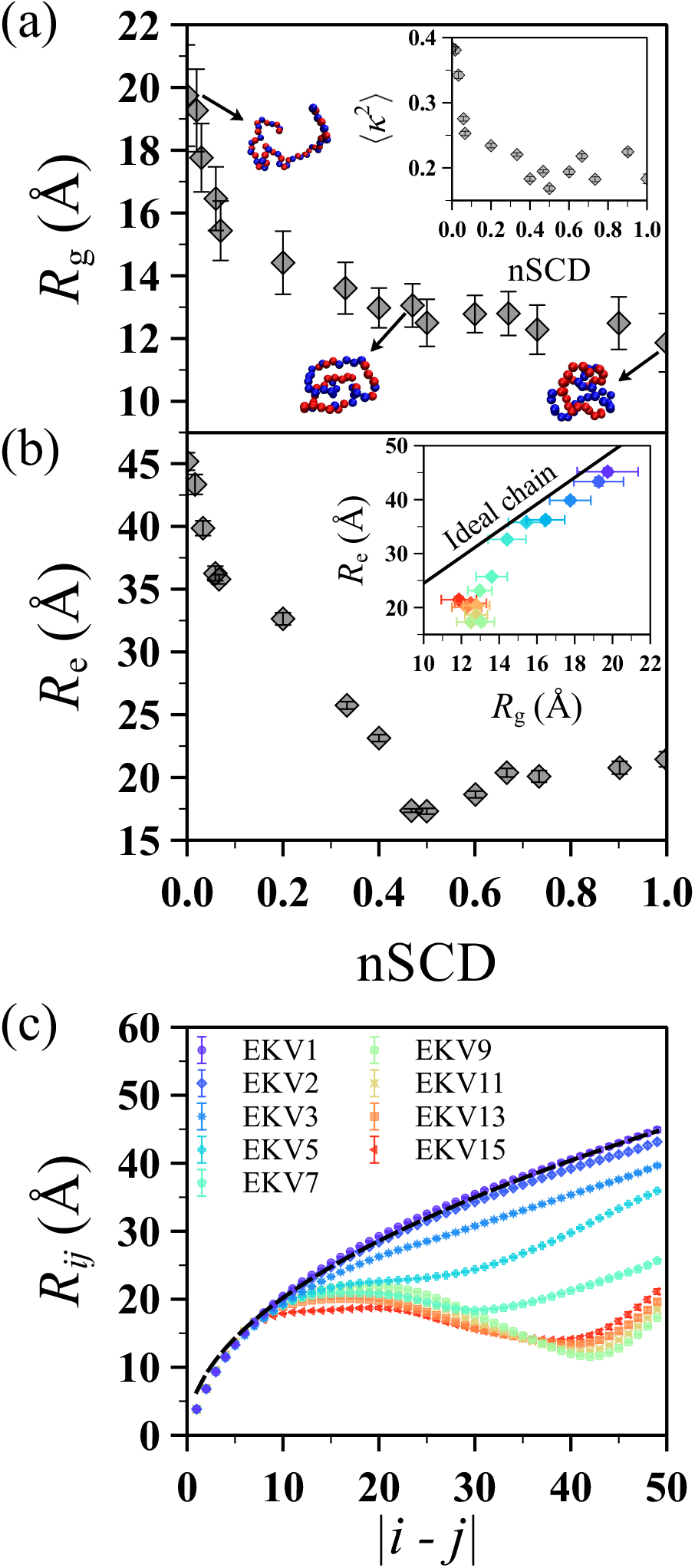
(a) Radius of gyration *R*_g_ as a function of nSCD for the EKVs in Figure 1 along with representative conformations for select EKVs. The inset shows the relative anisotropy ⟨*κ*^2^⟩ as a function of nSCD for the same EKVs. (b) End-to-end distance *R*_e_ as a function of nSCD for the same EKVs. The inset shows the correlation between *R*_g_ and *R*_e_ from the simulations compared to the theoretical expectation for an ideal chain 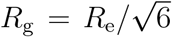. (c) Interresidue distance *R*_*ij*_ as a function of residue separation in the chain |*i* − *j*| for select EKVs. The dashed line corresponds to the ideal chain scaling *R*_*ij*_ = *b* |*i* − *j*|^1*/*2^, where *b* = 6.39 Å was fitted for EKV1 using the theoretically expected end-to-end distance 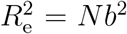 of an ideal chain with *N* = 50. The symbol color, ranging from purple to red, indicates increasing nSCD. The same color scale is used in the inset of (b).

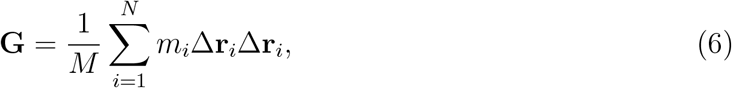

where Δ**r**_*i*_ is the vector to residue *i* from the chain’s center of mass and *M* is the total mass of the chain. We found that *R*_g_ monotonically decreased with increasing charge segregation (*i*.*e*., increasing nSCD) in a non-linear way: the most significant chain compaction occurred for nSCD ≲ 0.2, followed by a smaller decrease of *R*_g_ for larger nSCD. We also computed the probability distribution of *R*_g_ (Figure S2a) and found that it narrowed with increasing nSCD, indicating smaller conformational fluctuations for highly charge-segregated sequences. We further quantified the average shape of the EKVs using the average relative anisotropy ⟨*κ*^2^⟩ (inset of Figure 2a). We computed *κ*^2^ for a given chain conformation from the eigenvalues *λ*_*i*_ of **G**:^40^

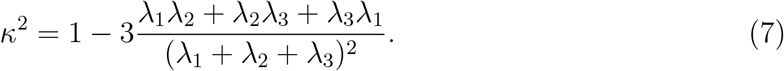

The values of ⟨*κ*^2^⟩ for a sphere and rod are 0 and 1, respectively. For small nSCD, the relative anisotropy was close to the numerically determined value for a three-dimensional random walk, ⟨*κ*^2^⟩ ≈ 0.39.^41^ Similar to *R*_g_, ⟨*κ*^2^⟩ also decreased with increasing nSCD with highly charge-segregated sequences showing a semi-compact conformation rather than a globular conformation as indicated by the non-zero ⟨*κ*^2^⟩ values. The dependence of *R*_g_ and ⟨*κ*^2^⟩ on nSCD for our model polyampholyte is consistent with previous theoretical and atomistic simulation studies of similar polyampholyte sequences. ^19,22^

Chain conformations can also be characterized through the root mean squared end-to-end distance, *R*_e_ = ⟨**R**_e_·**R**_e_⟩^1*/*2^,^42^ where **R**_e_ is the end-to-end vector pointing from the first residue to the last residue. Unlike *R*_g_, which showed a prominent decrease only until nSCD ≈ 0.2, *R*_e_ decreased strongly up to nSCD ≈ 0.5 (Figure 2b). For nSCD ≳ 0.5, we found little change (slight upward trend) in *R*_e_ with further increase in nSCD. We note that the same trends also persisted in the probability distribution of *R*_e_ (Figure S2b). As a consequence, within the statistical uncertainties, *R*_g_ and *R*_e_ for EKVs with nSCD ≲ 0.2 roughly correlated according to the theoretical expectation for an ideal chain 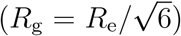^43^ but significantly deviated from this proportional scaling with increasing charge segregation (inset of Figure 2b). This suggests that *R*_e_ can be inferred reliably from *R*_g_ (and vice-versa) based on homopolymer models only for rather uniformly charge-patterned sequences. This finding is in line with previous experiments that observed decoupling between *R*_g_ and *R*_e_ for 10 naturally occurring heteropolymers,^44^ which leads to difficulties in interpreting their coil–globule transition.^45^

In our model polyampholyte, the heterogeneous interactions between residues are mainly due to electrostatics because the vdW interaction parameters for the E and K residues are similar. The effect of heterogeneity on the local chain conformations can be studied in more detail using the interresidue distance *R*_*ij*_ (Figure 2c).^19^ The average distance *R*_*ij*_ between two residues *i* and *j* was measured and then computed as a function of residue separation in the chain |*i* − *j*| by averaging over all pairs of residues in a given EKV. For sequences with nSCD ≲ 0.035 (EKV1 to EKV3), *R*_*ij*_ monotonically increased with increasing |*i* − *j*|, indicating only weak attraction between the (nearly) uniformly distributed E and K residues in the sequence. These sequences were found to closely follow the ideal chain scaling *R*_*ij*_ = *b*|*i* − *j*|^1*/*2^ with *b* being the segment size of the ideal chain. To illustrate this scaling behavior, we fit *b* = 6.39 Å for EKV1 using the end-to-end distance 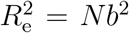 of an ideal chain, and plotted the expected *R*_*ij*_ as a dashed line in Figure 2c. For nSCD *>* 0.035, *R*_*ij*_ deviated strongly from ideal-chain scaling and even became non-monotonic. This behavior can be attributed to the electrostatic attractions between segregated E and K blocks in such sequences.

To further understand these trends in *R*_*ij*_, we computed interresidue distance-based contact maps and energy maps (Figures 3 and S3). For the distance-based contact maps, two residues *i* and *j* were considered to be in contact if the distance between them was less than 1.5 *σ*. The contact probability was computed as *P* = ⟨*n*_*ij*_⟩, where *n*_*ij*_ = 1 if the distance criteria for residues *i* and *j* is satisfied and *n*_*ij*_ = 0 if not. In the contact maps, the probabilities are plotted as − ln(*P/P*_max_), where *P*_max_ is the maximum probability observed between a pair of residues in a given sequence. For the energy maps, the pairwise non-bonded interaction energy *U*_nb_ = *U*_vdW_ + *U*_e_ was computed. Smaller values of − ln(*P/P*_max_) and *U*_nb_ correspond to a more favorable contact between a pair of residues. We excluded pairs of residues that were separated by one or two bonds from the map because their contacts were primarily determined by the chain connectivity. The purpose of showing both the distancebased contact maps and energy maps is that they are complementary but distinct metrics, and using both is especially helpful for reliably determining contacts between residues of different sizes.^46^ As seen from Figures 3 and S3, contacts were more evenly distributed along the chain contour for more uniformly charge-patterned sequences with nSCD ≲ 0.035 (EKV1 to EKV3). For nSCD *>* 0.035, contacts were localized at E and K residue blocks in such sequences. These observations indicate that the arrangement of charged residues in polyampholytes dictates the nature of intrachain molecular interactions (*i*.*e*., distributed interactions in low-nSCD EKVs *vs*. localized interactions in intermediate-and high-nSCD EKVs) that are well-described by the nSCD parameter.

**Figure 3:**
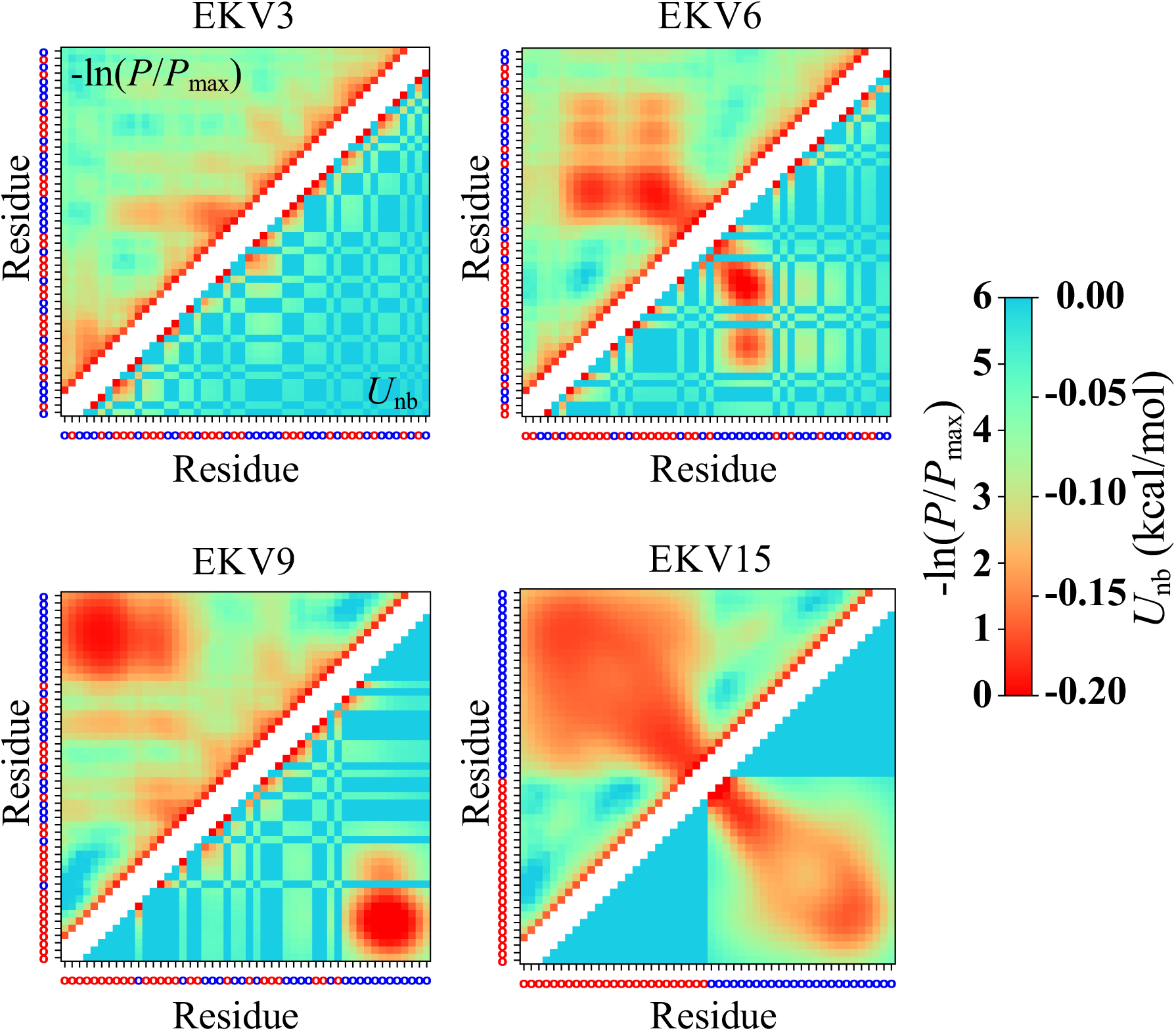
Interresidue distance-based contact map − ln(*P/P*_max_) (above diagonal) and energy map *U*_nb_ (below diagonal) for select EKVs. The residue at each position is shown as a red (E) or blue (K) circle, respectively, on the axes. Two diagonals on either side of the main diagonal are removed in the maps to exclude directly bonded residues and residues that are separated by two bonds.

### 3.2 Dynamics of model polyampholytes

Having characterized the effect of charge patterning on the conformational properties of our model polyampholyte, we then investigated the influence of charge patterning on its dynamical properties, which is an area that is far less understood. We first analyzed the end-to-end orientational dynamics that are representative of chain-level relaxation. Recently, the end-to-end motion of a single chain and of chains in the condensed phase was used as a proxy to infer the rheology of condensates formed by the low-complexity domain of fused in sarcoma (FUS).^47^ In experiments, the end-to-end dynamics of IDPs are often inferred from Förster resonance energy transfer experiments using dyes placed at the chain ends.^48^ In simulations, we can directly measure the end-to-end vector **R**_e_ and compute its autocorrelation function,

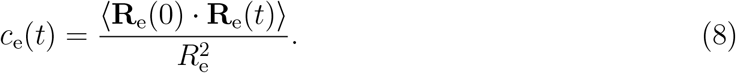

We found that *c*_e_ varied strongly between EKVs (Figure S4). To quantify this trend, we fit *c*_e_ to the Kohlrausch–Williams–Watts stretched exponential function *c*_e_(*t*) = exp[−(*t/τ*)^*β*^], with parameters *τ* and *β*,^49^ and then computed the corresponding relaxation time *τ*_e_ by integration,

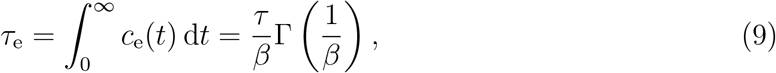

where Γ is the gamma function. To eliminate the chain length dependence of the end-to-end vector relaxation time, we normalize *τ*_e_ by 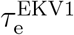, the relaxation time for EKV1 having nSCD = 0.

Interestingly, *τ*_e_ had a similar dependence on nSCD (Figure 4) as *R*_e_: it initially decreased significantly with increasing nSCD up until nSCD ≈ 0.5, but then changed only little (slight upward trend) for larger nSCD values. To assess the generality of this behavior, we performed additional simulations of both shorter (*N* = 30) and longer (*N* = 70) EKV chains (Figures S5 and S6). After normalizing *τ*_e_ by 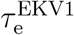 for each respective chain length, we found that the end-to-end orientational dynamics had a similar dependence on nSCD for all three chain lengths (Figure 4). To test the robustness of the nSCD parameter for classifying the different relaxation regimes, we also simulated sequences (*N* = 50) with similar nSCD values to EKV9 and EKV10 (nSCD ≈ 0.5) but different charge patterning near the chain ends (Figure S7). These sequences exhibited slightly different *τ*_e_ values when compared to the selected EKV9 and EKV10 (Figure 4), implying that the end-to-end interactions depend not only on the charge patterning of the overall sequence, but also on the local sequence details at the chain ends that are not captured by the nSCD parameter. Nevertheless, the *τ*_e_ values for these new sequences still fell close to the existing curve for the other EKVs with some scatter.

**Figure 4:**
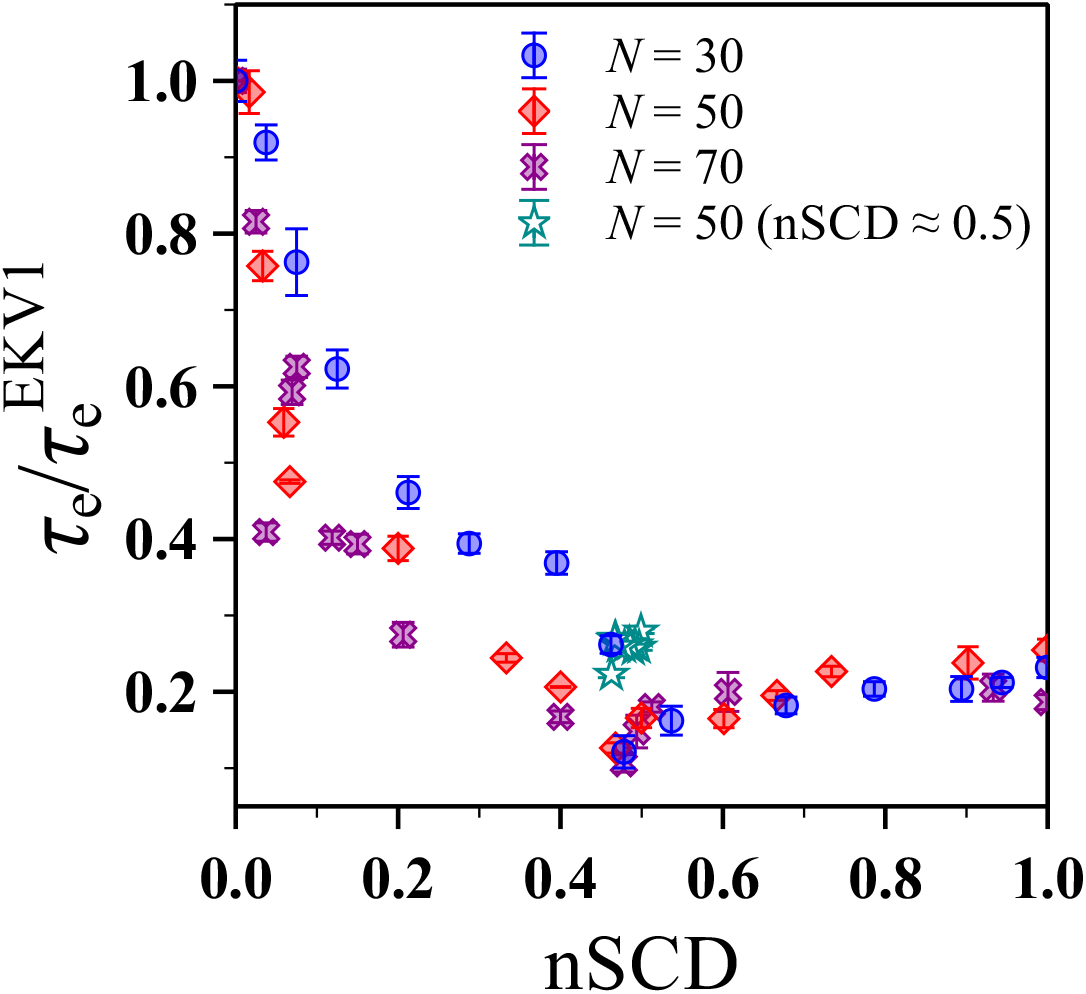
Relaxation time *τ*_e_ of the end-to-end vector, normalized by that of EKV1, as a function of nSCD for different chain lengths *N*.

Given that *R*_e_ and *τ*_e_ appeared to have a similar dependence on nSCD, we tested for correlation between the two by plotting 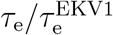 against 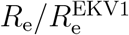 (Figure 5a). In this representation, the simulation data for different chain lengths fell on a master curve that followed a power law 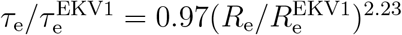, further corroborating the generality of the observed trends. We emphasize that this collapse was true even for those sequences with nSCD ≈ 0.5 (Figure S7) whose charge patterning at the chain ends was different than that of selected EKV9 and EKV10. To rationalize this behavior, we consider the Rouse model, ^50^ which is a simple description for the dynamics of an ideal (Gaussian) chain with free-draining hydrodynamics. There, the end-to-end vector relaxation time increases with increasing chain size as *τ*_e_ = *γN* ^2^*b*^2^*/*(3*π*^2^*k*_B_*T*), where *k*_B_ is the Boltzmann constant.^43^ Assuming that changes in *R*_e_ are due to changing *b* with fixed *N*, the normalized relaxation times should then follow 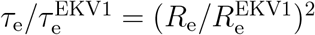 for an ideal chain homopolymer because *R*_e_ = *Nb*^2^. We emphasize, though, that this relationship does not necessarily hold for a heteropolymer that does not conform to ideal-chain statistics (Figure 2). The fact that the ideal-chain scaling of 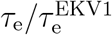 was similar to the power-law fit to the simulation data suggests that the EKVs have similar end-to-end dynamics as equivalent ideal chains with the same *N* and *R*_e_ but reduced segment length *b*. However, the fact that such rescaling works for features associated with the end-to-end vector does *not* mean that the EKVs follow ideal chain statistics at all length scales, as evidenced by the interresidue distances shown in Figure 2c.

**Figure 5:**
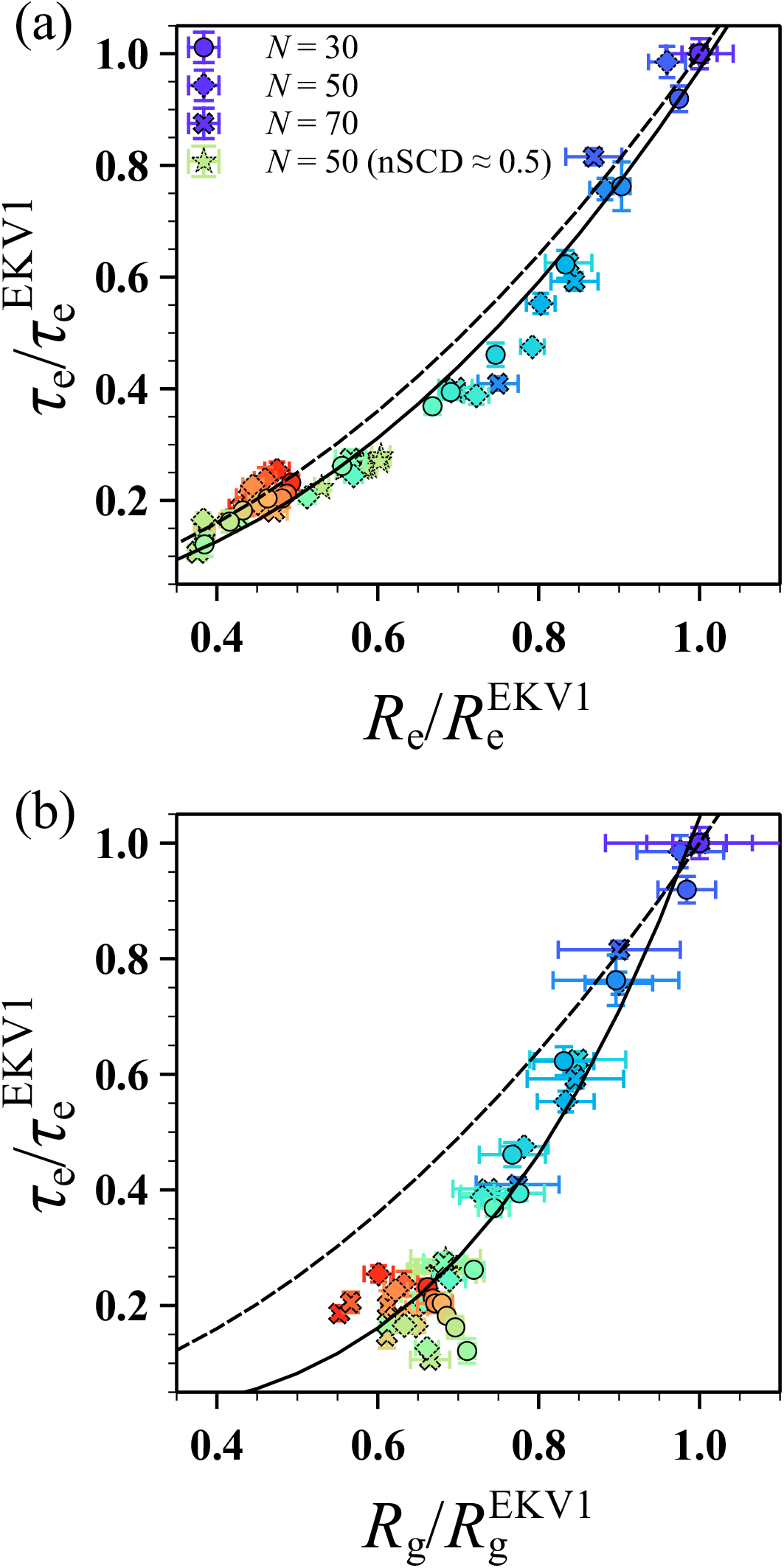
Relaxation time *τ*_e_ of the end-to-end vector as a function of (a) end-to-end distance *R*_e_ and (b) radius of gyration *R*_g_. Data shown for different chain lengths *N* and for sequences shown in Figure S7, with all values normalized by those of EKV1 for each *N*. The solid lines in (a) and (b) correspond to power law fits 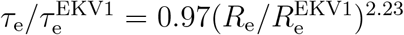 and 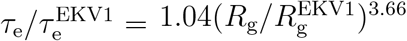, respectively. The dashed lines in (a) and (b) correspond to the Rouse model scaling for an ideal chain with fixed number of segments *N* and varying segment size *b*, which gives 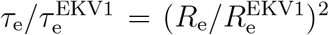 and 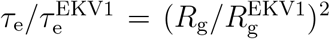. The symbol color, ranging from purple to red, indicates increasing nSCD.

This point was further highlighted when we plotted 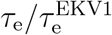 against 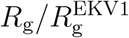 (Figure 5b). The data for the different chain lengths collapsed onto a master curve only for the low-nSCD sequences, following the power law 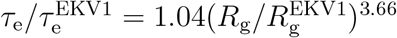, and was scattered for moderately to highly charge-segregated sequences. For an ideal chain, we would expect 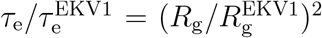 within the Rouse model because 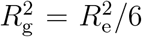, but only sequences with very low-nSCD values (nSCD ≲ 0.035) were close to this scaling. The different fitted scaling exponent and scattering of the data for more charge-segregated sequences is consistent with the observed decoupling between *R*_g_ and *R*_e_ (inset of Figure 2b). Indeed, the nSCD value above which significant deviation from ideal-chain scaling occurred was roughly the value at which *R*_*ij*_ also deviated (Figure 2c). Together, these findings indicate that the structural and dynamical properties associated with the end-to-end vector respond similarly to the increasing charge segregation in the sequences. This was true for EKVs both with low- and high-nSCD. However, the correlation between structural and dynamical properties is not straightforwardly captured by simple homopolymer models such as the Rouse model depending on the extent of charge segregration.

After investigating the chain-level dynamics, we considered the segmental dynamics associated with internal relaxation of the chain at different length scales. We computed the normal modes,^51–53^

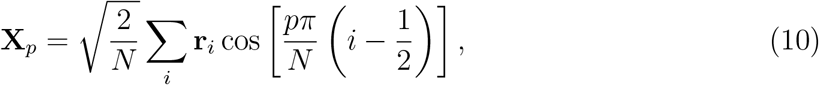

where 1 ≤ *p* ≤ *N* − 1. The autocorrelation function of **X**_*p*_,

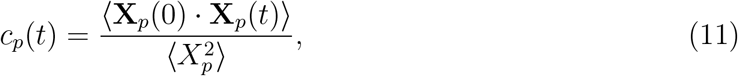

was used to compute the relaxation time *τ*_*p*_ of the *p*-th mode using the same procedure as we used for *c*_e_ and *τ*_e_ (Figure 6). Given that modes with larger *p* correspond to smaller segments of the chain, *τ*_*p*_ is expected to decrease monotonically with increasing *p*. For example, within the Rouse model,^51–54^

**Figure 6:**
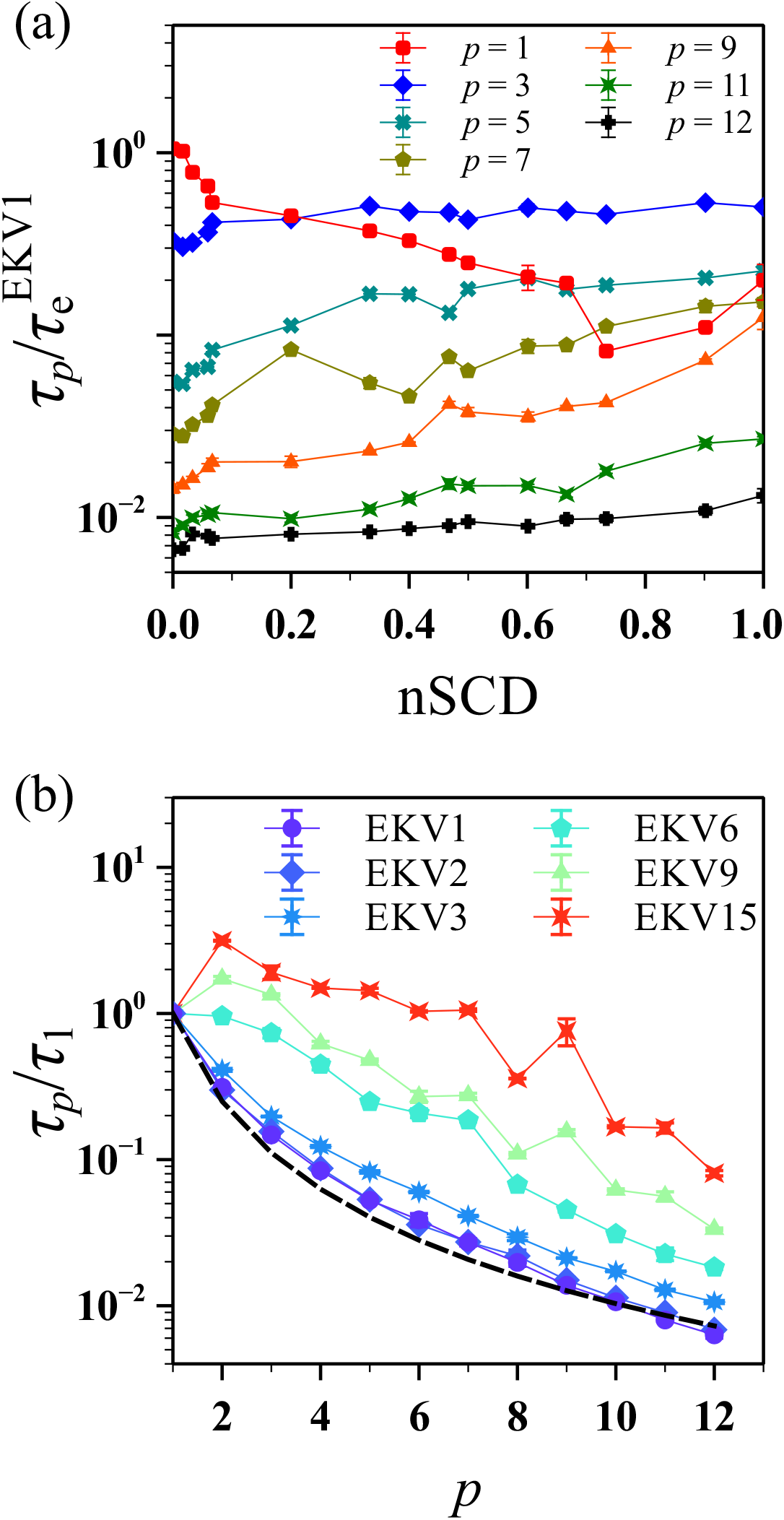
(a) Relaxation time *τ*_*p*_ of different normal modes, normalized by the end-to-end vector relaxation time of EKV1, as a function of nSCD for the EKVs. (b) Relaxation time *τ*_*p*_, normalized by that obtained for the *p* = 1 mode, as a function of mode index *p* for select EKVs. The dashed line corresponds to the theoretically expected scaling of the Rouse model for an ideal chain.

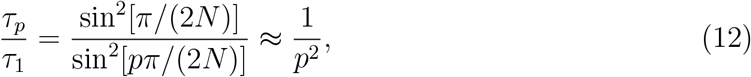

with the approximation holding only for small *p*. However, we observed a monotonic decrease in *τ*_*p*_ with *p* only for EKVs with nSCD ≲ 0.2 (Figure 6a). For moderately to highly charge-segregated sequences (nSCD *>* 0.2), the *p* = 1 relaxation mode became significantly slower than the higher (*p >* 1) modes, indicating the formation of slowly changing structures at intermediate length-scales (see Figures 2 and 3). To highlight this behavior, we plotted *τ*_*p*_*/τ*_1_ as a function of *p* for select EKVs and compared them to the Rouse model prediction for an ideal chain (Figure 6b).^43^ Consistent with *τ*_e_ (Figure 5b), the more uniformly charge-patterned sequences (EKV1 to EKV3 whose nSCD ≲ 0.035) closely followed the Rouse scaling, but the more charge-segregated EKVs deviated significantly from it. Thus, our findings suggest that the dynamics exhibited by (nearly) uniformly patterned polyampholytes in solution might be interpreted using simple homopolymer scaling laws. In contrast, the same scaling laws should be used with caution for polyampholytic IDPs with moderate to high charge segregation.

Having assessed both the chain end-to-end and segmental dynamics, we last explored the fine-scale dynamics associated with the relaxation of individual bonds. In experiments, the backbone amide bond vector is typically used to compute the residue-wise rigidity for identifying regions in IDPs that could play a major role in the overall protein dynamics.^55^ It is thus important to understand the bond relaxation in EKVs with different degrees of charge segregation. Given that only the low-nSCD sequences followed typical homopolymer scaling predictions for the structural and dynamical properties studied so far, we hypothesized that the relaxation of the bonds away from the chain ends in such low-nSCD sequences would be similar. To test this hypothesis, we computed the autocorrelation function *c*_b_ and associated relaxation time *τ*_b_ for each of the 49 bond vectors in a given EKV using the same methodology as for the end-to-end vector (*c*_e_ and *τ*_e_). We then normalized *τ*_b_ for each bond by the average relaxation time of all bonds in EKV1 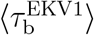 (Figure 7). The bonds near the termini relaxed much faster than the ones near the center of the chain, as expected because of chain end effects. Most importantly, central bonds in the low-nSCD sequences (EKV1 and EKV2) relaxed in a similar fashion, which is seen in the near-constant profile of the relaxation times as a function of bond index. The bond relaxation times of EKV3 showed some variation with bond index when compared to the profiles of EKV1 and EKV2, indicating more pronounced electrostatic interactions in certain regions of the sequence. EKV9 showed a distinct slowing of the bond relaxation near the chain termini while bonds near the chain center behaved similar to the EKV1 sequence. This is likely due to the fact that EKV9 (nSCD close to 0.5) had blocks of oppositely charged E and K residues at the chain ends but a more uniform patterning in the middle of the chain. This effect was even more pronounced for the highly charge-segregated diblock EKV15 (nSCD = 1). These observations validated our hypothesis that all the bonds away from the termini in the more uniformly charge-patterned sequences would relax similarly, whereas those in the charge-segregated sequences would relax differently because of pronounced electrostatic interactions that depend on the degree of blockiness within them.

**Figure 7:**
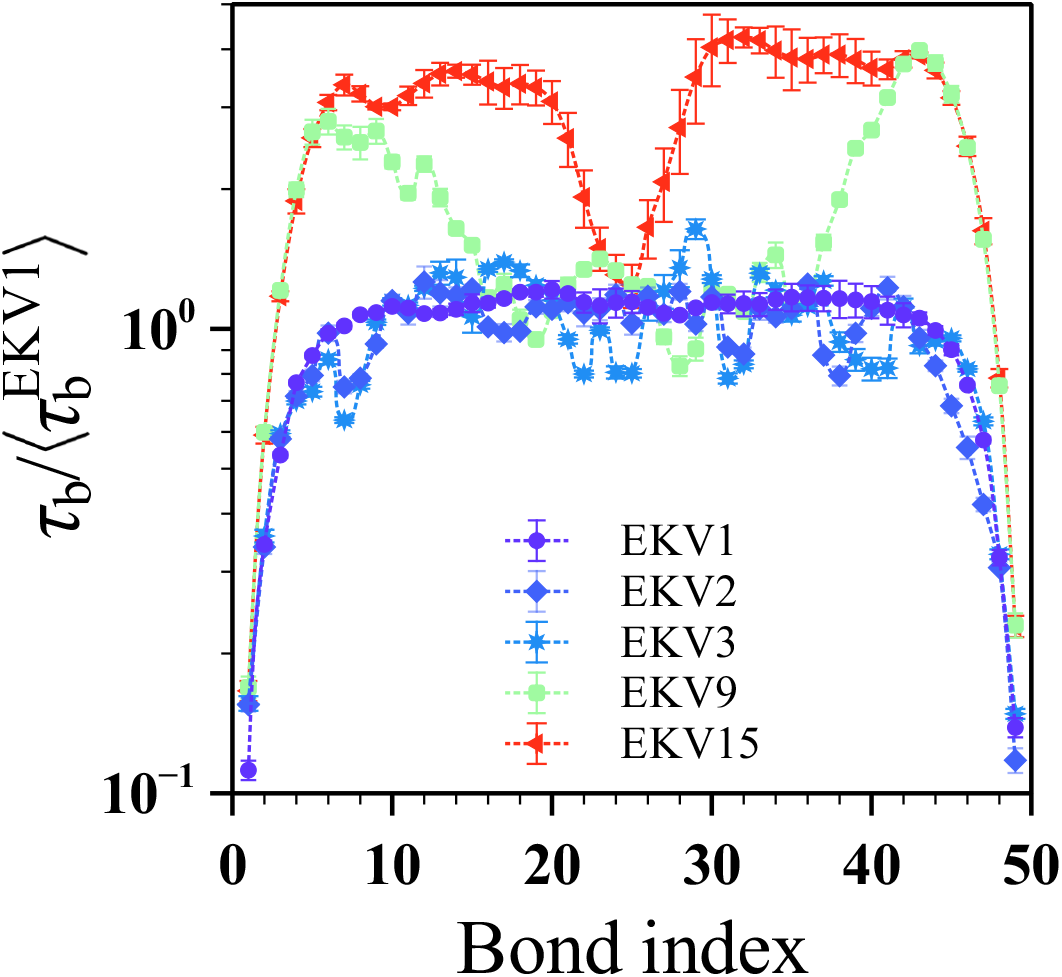
Relaxation time *τ*_b_ of the bond vectors, normalized by the average relaxation time of all bonds in EKV1, as a function of bond index for select EKVs.

### 3.3 Dynamics of naturally occurring polyampholytes

Our simulations of the model polyampholytic IDP revealed that its conformations and dynamics strongly depended on the sequence charge patterning. We then asked if these behaviors would also hold for naturally occurring polyampholytic IDPs. To this end, we investigated the segmental dynamics of a charge-rich IDR (*N* = 176) of the naturally occurring protein LAF-1. We focused on the wild-type sequence LAF-1 RGG WT and its shuffled variant LAF-1 RGG SHUF (Figure 8a), which were recently used to demonstrate that charge segregation increases the propensity of IDP solutions to form a condensed phase. ^27^ In that study, the LAF-1 RGG SHUF sequence was created by shuffling the residues in the LAF-1 RGG WT sequence such that the charges are more segregated. The uncharged residues in these sequences interacted with all the other residues through the van der Waals interactions as given in Equations 3 and 4. The LAF-1 RGG WT sequence is (nearly) uniformly charge-patterned as indicated by its high SCD value of -0.277, whereas LAF-1 RGG SHUF is moderately charge-segregated with SCD = -5.021. To put these SCD values in perspective, we determined the maximum and minimum SCD values achievable for sequences of the same length and composition as LAF-1 RGG, finding -0.003 and -28.183, respectively. Using these bounds, we determined the nSCD values for LAF-1 RGG WT and LAF-1 RGG SHUF as 0.010 and 0.178, respectively.

**Figure 8:**
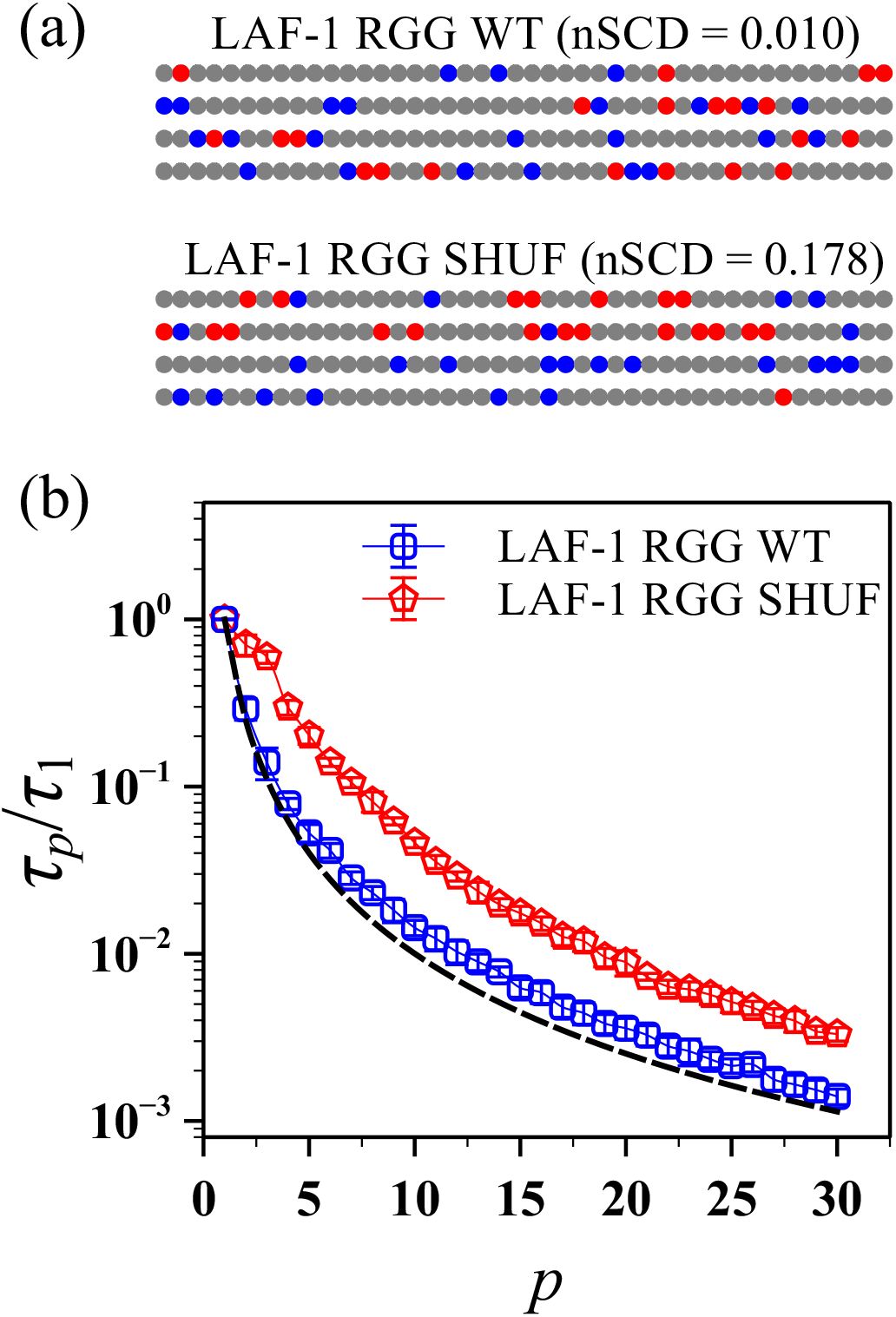
(a) Amino acid sequences of LAF-1 RGG WT and its shuffled charge-segregated variant LAF-1 RGG SHUF, each of length *N* = 176 residues. Negatively charged, positively charged, and uncharged residues are shown as red, blue, and gray circles, respectively. (b) Relaxation time *τ*_*p*_, normalized by that obtained for the *p* = 1 mode, as a function of mode index *p* for LAF-1 RGG WT and LAF-1 RGG SHUF sequences. The dashed line corresponds to the theoretically expected scaling of the Rouse model for an ideal chain.

Based on these nSCD values, the LAF-1 RGG WT and LAF-1 RGG SHUF sequences can be considered equivalent to EKV2 and EKV6 (Figure 1). Thus, we expected that LAF-1 RGG WT would exhibit dynamics following the Rouse model, whereas LAF-1 RGG SHUF should show some deviations. As seen in Figure 8b, the normalized relaxation time *τ*_*p*_*/τ*_1_ of LAF-1 RGG WT indeed closely followed the Rouse scaling for an ideal chain, whereas *τ*_*p*_*/τ*_1_ of LAF-1 RGG SHUF deviated from the theoretical prediction. We also note that unlike EKV6, which consisted entirely of charged residues, *τ*_*p*_*/τ*_1_ of LAF-1 RGG SHUF, which consisted of both uncharged and charged residues, decreased monotonically with increasing *p*.

### 3.4 Effect of hydrodynamic interactions on polyampholyte dynamics

The results presented so far were obtained from LD simulations that did not account for solvent-mediated hydrodynamic interactions (HI) between residues. In dilute solutions, however, HI have a strong impact on chain dynamics,^54,56^ and it is important to verify that the general trends in dynamical properties found in Sections 3.2 and 3.3 for the model polyampholyte and LAF-1 RGG variants, respectively, hold when HI are taken into account. For this purpose, we performed additional simulations using the multiparticle collision dynamics (MPCD) technique.^57–59^ MPCD uses a particle-based, mesoscopic solvent that undergoes alternating streaming and collision steps to propagate HI. During the streaming step of our simulations, the solvent particles moved ballistically over a time interval 100 fs. The residues were simultaneously integrated forward in time over this interval using the standard velocity-Verlet algorithm with the same 10 fs timestep as in the LD simulations. ^60^ The solvent particles and residues then participated in a linear-momentum-conserving collision in a spatially localized cubic cell (edge length *a* = 3.8 Å) according to the stochastic rotation dynamics rule.^57^ A rotation angle of 130 ° was used for the collisions, and the cubic cells that contained the solute and the solvent particles were randomly shifted to preserve Galilean invariance.^61^ The mass of a solvent particle was set to 25 g*/*mol and the number density of the solvent particles was fixed at 5 */a*^3^ (0.091*/* Å^3^). These values ensured that the solvent mass in a cubic cell was approximately the same as the residue mass.^62^ A constant temperature of 300 K was maintained using a cell-level Maxwellian thermostat. ^63^ The chosen values for the MPCD solvent result in a dynamic viscosity 0.42 cP and Schmidt number Sc ≈ 18, which indicates that the solvent is liquid-like. For these MPCD simulations, the edge length of the cubic simulation box was chosen to be 159.6 Å to give an integer number of collision cells in all directions. The simulations were also carried out using HOOMD-blue (version 2.9.3). ^64^

The chain-level dynamics, as quantified by the end-to-end vector relaxation time *τ*_e_, were slower in the MPCD simulations (+HI) than in the LD simulations (−HI) for all EKVs (Figure S8). This quantitative difference was because of the higher solvent viscosity in MPCD than that inferred in LD simulations. However, *τ*_e_ from the +HI simulations collapsed onto a similar curve as a function of nSCD as the −HI simulations (Figure 9a) after we normalized it by 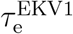. We then computed the normal modes and the segmental relaxation in the +HI simulations (Figure 9b). The Rouse model is not appropriate for describing an ideal chain with HI; instead, we used the Zimm model^56^ as a theoretical baseline for *τ*_*p*_,^65,66^

**Figure 9:**
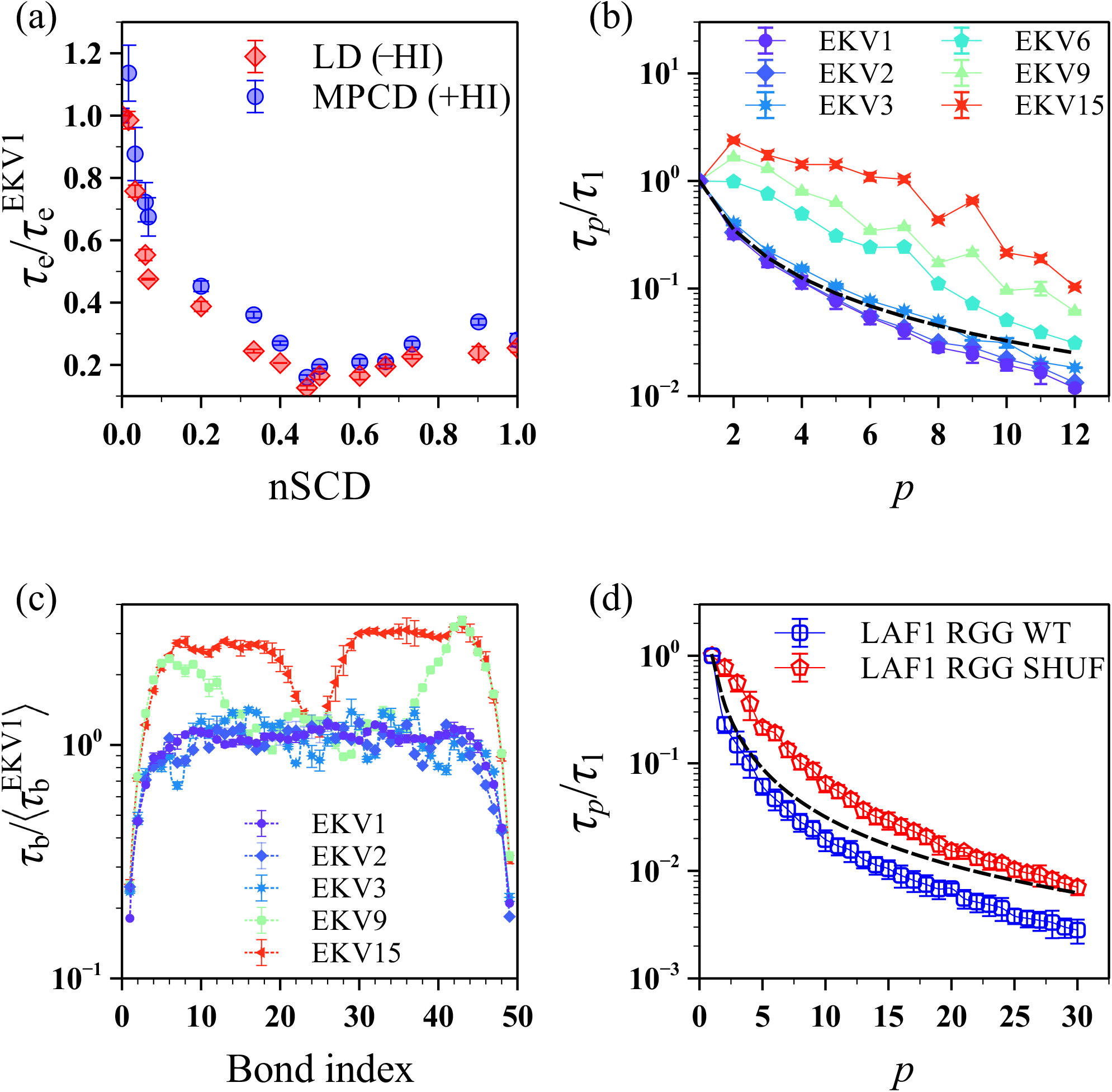
Relaxation times of the (a) end-to-end vector *τ*_e_, (b) normal modes *τ*_*p*_, and (c) bond vectors *τ*_b_ for the model polyampholyte EKVs from the MPCD (+HI) and LD (−HI) simulations. (d) Relaxation time *τ*_*p*_ of the normal modes for LAF-1 RGG WT and LAF-1 RGG SHUF from the MPCD (+HI) simulations. All quantities are normalized as in Figures 4, 6, 7, and 8. The dashed lines in (b) and (d) are the expected scaling of the Zimm model for an ideal chain.

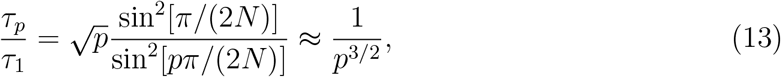

with the approximation holding only for small *p*. We found that the relaxation times of modes with smaller *p* for the more uniformly charge-patterned sequences (EKV1 to EKV3 whose nSCD ≲ 0.035) closely followed the Zimm scaling but deviated with increasing *p* (Figure 9b). In contrast, for the moderately to highly charge-segregated sequences, all normal modes failed to follow the Zimm scaling. Last, analogous to the −HI simulations (Figure 7), the fine-scale dynamics as quantified by the local bond relaxation exhibited a near-constant profile for the bonds away from the chain termini in the more uniformly charge-patterned sequences (Figure 9c).

When considering LAF-1 RGG WT, only the relaxation times of modes with *p* ≲ 7 closely followed the Zimm scaling. For the more charge-segregated LAF-1 RGG SHUF, the modes decayed much more slowly (Figure 9d). Although there are some (expected) quantitative differences between the chain dynamics in the simulations with and without HI, we observed in both cases the same general trends for the chain dynamics when increasing the charge segregation in the EKV and LAF-1 RGG sequences. This suggests that the differing behavior of polyampholytic IDPs arises primarily from altered intrachain molecular interactions as a result of sequence charge patterning, independent of the presence of hydrodynamic interactions.

## 4 Conclusions

In this work, we have elucidated the effect of charge patterning on the conformation as well as the chain-level and segmental dynamics of polyampholytes in solution, with a focus on model polyampholytes consisting of glutamic acid (E) and lysine (K) residues with zero net charge. We performed coarse-grained simulations (single bead per residue) with and without hydrodynamic interactions, and quantified the sequence charge patterning in the E–K variants (EKVs) *via* the normalized sequence charge decoration (nSCD) parameter that goes from 0 (perfectly alternating sequence) to 1 (diblock sequence). With increasing charge segregation, we observed a transition from ideal chain-like to semi-compact polymer conformations, which we quantified through the radius of gyration *R*_g_, the end-to-end distance *R*_e_, and the relative shape anisotropy *κ*^2^. Though both *R*_g_ and *R*_e_ decreased with increasing nSCD, the rate of decrease was prominent only until nSCD ≈ 0.2 for *R*_g_ and nSCD ≈ 0.5 for *R*_e_. Consequently, the low-nSCD EKVs (nSCD ≲ 0.2) roughly followed the theoretically expected ideal chain correlation 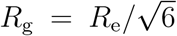, whereas the high-nSCD EKVs deviated strongly, thus highlighting the caveats in interpreting the size measures of charge-segregated sequences using simple homopolymer models. This finding was further supported by observing that the interresidue distance *R*_*ij*_ of only low-nSCD EKVs followed the ideal chain scaling ∼ *N* ^1*/*2^. To better understand this behavior, we computed distance- and energy-based interresidue contact maps, which revealed the formation of non-covalent bonds between blocks of oppositely charged residues for the intermediate- and high-nSCD EKVs.

To study the effect of these structural changes on the chain dynamics, we characterized the chain-level dynamics by computing the relaxation time *τ*_e_ of the end-to-end vector, finding that *τ*_e_ decreased with increasing nSCD, similar as *R*_e_. Specifically, *τ*_e_ and *R*_e_ values for different chain lengths *N*, when normalized by the corresponding values of the perfectly alternating sequence with nSCD = 0, collapsed onto a master curve for the entire range of nSCD. We rationalized this behavior by considering the end-to-end dynamics of a Rouse chain with the same *R*_e_ and *N* as the EKVs, which showed the same scaling behavior across the entire nSCD range. Two important conclusions emerge from these findings: (1) the conformation and dynamics associated with the end-to-end vector of polyampholytes show similar correlation with nSCD, and (2) the extent of charge segregation dictates the applicability of simple homopolymer models to describe the structure and dynamics of polyampholytes. These findings were further substantiated by an analysis of the segmental dynamics, which revealed that the relaxation time of the normal modes closely followed the theoretically expected scaling of the Rouse model for an ideal chain (Zimm model when we included hydrodynamic interactions) only for the more uniformly charged-patterned sequences. This behavior probably originates from the negligible net electrostatic attractions in low-nSCD EKVs, as indicated by our contact analysis and the near-constant values of local bond relaxation time away from the termini in such sequences.

To demonstrate the pertinence of our findings to naturally occurring disordered regions with both charged and uncharged residues, we also studied the structure and dynamics of LAF-1 RGG variants. In line with our results for the model polyampholytes, the normal modes of the uniformly charge-patterned protein LAF-1 RGG WT followed classic homopolymer models, whereas the charge-segregated variant LAF-1 RGG SHUF did not. These results highlight the effect of charge distribution on the single-chain polyampholytic IDP dynamics that serve as a first step towards investigating their dynamics and rheology in dilute and condensed phases. We believe that a detailed understanding of IDP dynamics and rheology in dilute and condensed phases could help in establishing structure–dynamics–rheology– function relationships for IDPs, thus facilitating the design of protein sequences with desired dynamical responses and material properties.

## Acknowledgments

Research reported in this work was supported by the National Institute of General Medical Science of the National Institutes of Health under the grant R01GM136917 and the Welch Foundation under the grant A-2113-20220331. A.N. acknowledges funding by the Deutsche Forschungsgemeinschaft (DFG, German Research Foundation) through projects 274340645 and 470113688. We also gratefully acknowledge the Texas A&M High Performance Research Computing (HPRC) for providing the computational resources to carry out the simulations discussed in this work.

## TOC Graphic

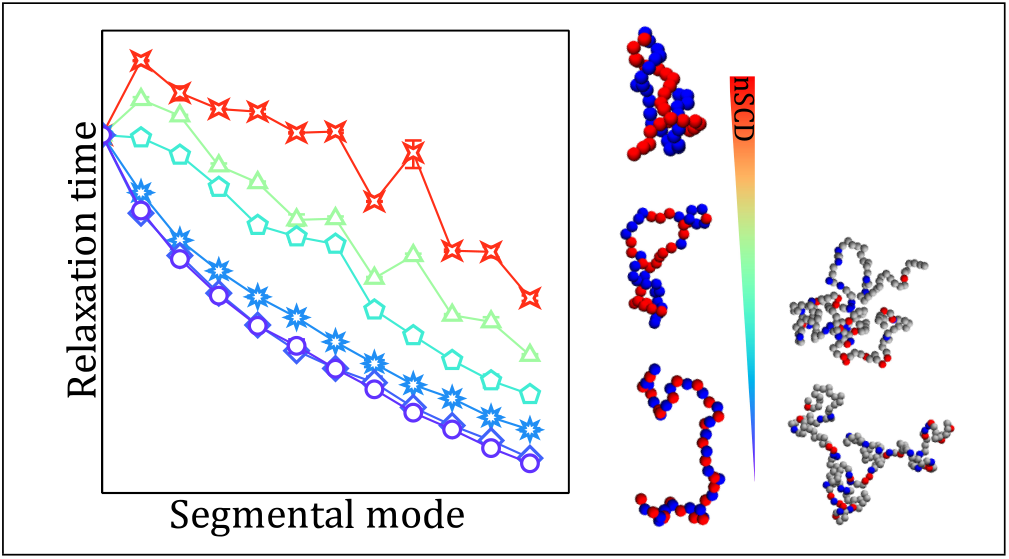

## References

(1) Brangwynne, C. P.; Eckmann, C. R.; Courson, D. S.; Rybarska, A.; Hoege, C.; Gharakhani, J.; Jülicher, F.; Hyman, A. A. Germline P Granules Are Liquid Droplets That Localize by Controlled Dissolution/Condensation. Science 2009, 324, 1729–1732.

(2) Dignon, G. L.; Best, R. B.; Mittal, J. Biomolecular Phase Separation: From Molecular Driving Forces to Macroscopic Properties. Annual Review of Physical Chemistry 2020, 71, 53–75.

(3) Hammer, S. K.; Avalos, J. L. Harnessing Yeast Organelles for Metabolic Engineering. Nature Chemical Biology 2017, 13, 823–832.

(4) Kerfeld, C. A.; Aussignargues, C.; Zarzycki, J.; Cai, F.; Sutter, M. Bacterial Micro-compartments. Nature Reviews Microbiology 2018, 16, 277–290.

(5) Alberti, S.; Gladfelter, A.; Mittag, T. Considerations and Challenges in Studying Liquid-Liquid Phase Separation and Biomolecular Condensates. Cell 2019, 176, 419– 434.

(6) Sawyer, I. A.; Bartek, J.; Dundr, M. Phase Separated Microenvironments Inside the Cell Nucleus are Linked to Disease and Regulate Epigenetic State, Transcription and RNA Processing. Seminars in Cell & Developmental Biology 2019, 90, 94–103, 3D Genome and Diseases.

(7) Forman-Kay, J. D.; Kriwacki, R. W.; Seydoux, G. Phase Separation in Biology and Disease. Journal of Molecular Biology 2018, 430, 4603–4606.

(8) Sickmeier, M.; Hamilton, J. A.; LeGall, T.; Vacic, V.; Cortese, M. S.; Tantos, A.; Szabo, B.; Tompa, P.; Chen, J.; Uversky, V. N. et al. DisProt: The Database of Disordered Proteins. Nucleic Acids Research 2006, 35, D786–D793.

(9) Madinya, J. J.; Chang, L.-W.; Perry, S. L.; Sing, C. E. Sequence-dependent self-coacervation in high charge-density polyampholytes. Mol. Syst. Des. Eng. 2020, 5, 632–644.

(10) Srivastava, D.; Muthukumar, M. Sequence Dependence of Conformations of Polyam-pholytes. Macromolecules 1996, 29, 2324–2326.

(11) Yamakov, V.; Milchev, A.; Jörg Limbach, H.; Dünweg, B.; Everaers, R. Conformations of Random Polyampholytes. Phys. Rev. Lett. 2000, 85, 4305–4308.

(12) Higgs, P. G.; Joanny, J. Theory of Polyampholyte Solutions. The Journal of Chemical Physics 1991, 94, 1543–1554.

(13) Gutin, A. M.; Shakhnovich, E. I. Effect of a Net Charge on the Conformation of Polyam-pholytes. Phys. Rev. E 1994, 50, R3322–R3325.

(14) Edwards, S. F.; King, P. R.; Pincus, P. Phase Changes in Polyampholytes. Ferroelectrics 1980, 30, 3–6.

(15) Dobrynin, A. V.; Colby, R. H.; Rubinstein, M. Polyampholytes. Journal of Polymer Science Part B: Polymer Physics 2004, 42, 3513–3538.

(16) Wang, Z.; Rubinstein, M. Regimes of Conformational Transitions of a Diblock Polyampholyte. Macromolecules 2006, 39, 5897–5912.

(17) Samanta, H. S.; Chakraborty, D.; Thirumalai, D. Charge Fluctuation Effects on the Shape of Flexible Polyampholytes with Applications to Intrinsically Disordered Proteins. The Journal of Chemical Physics 2018, 149, 163323.

(18) Rumyantsev, A. M.; Johner, A.; de Pablo, J. J. Sequence Blockiness Controls the Structure of Polyampholyte Necklaces. ACS Macro Letters 2021, 10, 1048–1054.

(19) Das, R. K.; Pappu, R. V. Conformations of Intrinsically Disordered Proteins are Influenced by Linear Sequence Distributions of Oppositely Charged Residues. Proceedings of the National Academy of Sciences 2013, 110, 13392–13397.

(20) Bianchi, G.; Longhi, S.; Grandori, R.; Brocca, S. Relevance of Electrostatic Charges in Compactness, Aggregation, and Phase Separation of Intrinsically Disordered Proteins. International Journal of Molecular Sciences 2020, 21.

(21) Mao, A. H.; Crick, S. L.; Vitalis, A.; Chicoine, C. L.; Pappu, R. V. Net charge per residue modulates conformational ensembles of intrinsically disordered proteins. Proceedings of the National Academy of Sciences 2010, 107, 8183–8188.

(22) Sawle, L.; Ghosh, K. A Theoretical Method to Compute Sequence Dependent Configurational Properties in Charged Polymers and Proteins. The Journal of Chemical Physics 2015, 143, 085101.

(23) Firman, T.; Ghosh, K. Sequence Charge Decoration Dictates Coil-Globule Transition in Intrinsically Disordered Proteins. The Journal of Chemical Physics 2018, 148, 123305.

(24) Lin, Y.-H.; Chan, H. S. Phase Separation and Single-Chain Compactness of Charged Disordered Proteins Are Strongly Correlated. Biophysical Journal 2017, 112, 2043– 2046.

(25) Dignon, G. L.; Zheng, W.; Best, R. B.; Kim, Y. C.; Mittal, J. Relation between singlemolecule properties and phase behavior of intrinsically disordered proteins. Proceedings of the National Academy of Sciences 2018, 115, 9929–9934.

(26) Hazra, M. K.; Levy, Y. Charge Pattern Affects the Structure and Dynamics of Polyampholyte Condensates. Phys. Chem. Chem. Phys. 2020, 22, 19368–19375.

(27) Schuster, B. S.; Dignon, G. L.; Tang, W. S.; Kelley, F. M.; Ranganath, A. K.; Jahnke, C. N.; Simpkins, A. G.; Regy, R. M.; Hammer, D. A.; Good, M. C. et al. Identifying Sequence Perturbations to an Intrinsically Disordered Protein that Determine its Phase-Separation Behavior. Proceedings of the National Academy of Sciences 2020, 117, 11421–11431.

(28) Regy, R. M.; Dignon, G. L.; Zheng, W.; Kim, Y. C.; Mittal, J. Sequence dependent phase separation of protein-polynucleotide mixtures elucidated using molecular simulations. Nucleic Acids Research 2020, 48, 12593–12603.

(29) Baul, U.; Chakraborty, D.; Mugnai, M. L.; Straub, J. E.; Thirumalai, D. Sequence Effects on Size, Shape, and Structural Heterogeneity in Intrinsically Disordered Proteins. The Journal of Physical Chemistry B 2019, 123, 3462–3474.

(30) Rana, U.; Brangwynne, C. P.; Panagiotopoulos, A. Z. Phase separation vs aggregation behavior for model disordered proteins. The Journal of Chemical Physics 2021, 155, 125101.

(31) Perdikari, T. M.; Jovic, N.; Dignon, G. L.; Kim, Y. C.; Fawzi, N. L.; Mittal, J. A Predictive Coarse-Grained Model for Position-Specific Effects of Post-Translational Modifications. Biophysical Journal 2021, 120, 1187–1197.

(32) Ashbaugh, H. S.; Hatch, H. W. Natively Unfolded Protein Stability as a Coil-to-Globule Transition in Charge/Hydropathy Space. Journal of the American Chemical Society 2008, 130, 9536–9542.

(33) Miller, C. M.; Kim, Y. C.; Mittal, J. Protein Composition Determines the Effect of Crowding on the Properties of Disordered Proteins. Biophysical journal 2016, 111, 28–37.

(34) Weeks, J. D.; Chandler, D.; Andersen, H. C. Role of Repulsive Forces in Determining the Equilibrium Structure of Simple Liquids. The Journal of Chemical Physics 1971, 54, 5237–5247.

(35) Dignon, G. L.; Zheng, W.; Kim, Y. C.; Best, R. B.; Mittal, J. Sequence Determinants of Protein Phase Behavior from a Coarse-Grained Model. PLOS Computational Biology 2018, 14, 1–23.

(36) Kapcha, L. H.; Rossky, P. J. A Simple Atomic-Level Hydrophobicity Scale Reveals Protein Interfacial Structure. Journal of Molecular Biology 2014, 426, 484–498.

(37) Debye, P.; Hückel, E. De La Theorie Des Electrolytes. I. Abaissement Du Point De Congelation Et Phenomenes Associes. Physikalische Zeitschrift 1923, 24, 185–206.

(38) Anderson, J. A.; Glaser, J.; Glotzer, S. C. HOOMD-blue: A Python Package for High-Performance Molecular Dynamics and Hard Particle Monte Carlo Simulations. Computational Materials Science 2020, 173, 109363.

(39) https://github.com/mphowardlab/azplugins.

(40) Khabaz, F.; Khare, R. Effect of Chain Architecture on the Size, Shape, and Intrinsic Viscosity of Chains in Polymer Solutions: A Molecular Simulation Study. The Journal of Chemical Physics 2014, 141, 214904.

(41) Rudnick, J.; Gaspari, G. The Shapes of Random Walks. Science 1987, 237, 384–389.

(42) Flory, P. J. Principles of Polymer Chemistry; Cornell University Press, Ithaca, NY, 1953.

(43) Rubinstein, M.; Colby, R. H., et al. Polymer Physics; Oxford University Press, New York, 2003; Vol. 23.

(44) Fuertes, G.; Banterle, N.; Ruff, K. M.; Chowdhury, A.; Mercadante, D.; Koehler, C.; Kachala, M.; Estrada Girona, G.; Milles, S.; Mishra, A. et al. Decoupling of size and shape fluctuations in heteropolymeric sequences reconciles discrepancies in SAXS vs. FRET measurements. Proceedings of the National Academy of Sciences 2017, 114, E6342–E6351.

(45) Das, R. K.; Ruff, K. M.; Pappu, R. V. Relating sequence encoded information to form and function of intrinsically disordered proteins. Current Opinion in Structural Biology 2015, 32, 102–112.

(46) Bauer, D. J.; Stelzl, L. S.; Nikoubashman, A. Single-chain and condensed-state behavior of hnRNPA1 from molecular simulations. bioRxiv 2022,

(47) Benayad, Z.; von Bülow, S.; Stelzl, L. S.; Hummer, G. Simulation of FUS Protein Condensates with an Adapted Coarse-Grained Model. Journal of Chemical Theory and Computation 2021, 17, 525–537.

(48) Schuler, B.; Soranno, A.; Hofmann, H.; Nettels, D. Single-Molecule FRET Spectroscopy and the Polymer Physics of Unfolded and Intrinsically Disordered Proteins. Annual Review of Biophysics 2016, 45, 207–231.

(49) Larson, R. The Structure and Rheology of Complex Fluids; Oxford University Press, New York, 1998.

(50) Rouse, P. E. A Theory of the Linear Viscoelastic Properties of Dilute Solutions of Coiling Polymers. The Journal of Chemical Physics 1953, 21, 1272–1280.

(51) Kopf, A.; Dünweg, B.; Paul, W. Dynamics of polymer “isotope” mixtures: Molecular dynamics simulation and Rouse model analysis. The Journal of Chemical Physics 1997, 107, 6945–6955.

(52) Kalathi, J. T.; Kumar, S. K.; Rubinstein, M.; Grest, G. S. Rouse mode analysis of chain relaxation in polymer nanocomposites. Soft Matter 2015, 11, 4123–4132.

(53) Kalathi, J. T.; Kumar, S. K.; Rubinstein, M.; Grest, G. S. Rouse Mode Analysis of Chain Relaxation in Homopolymer Melts. Macromolecules 2014, 47, 6925–6931.

(54) Nikoubashman, A.; Milchev, A.; Binder, K. Dynamics of single semiflexible polymers in dilute solution. The Journal of Chemical Physics 2016, 145, 234903.

(55) Maiti, S.; De, S. Identification of Potential Short Linear Motifs (SLiMs) in Intrinsically Disordered Sequences of Proteins by Fast Time-Scale Backbone Dynamics. Journal of Magnetic Resonance Open 2022, 10-11, 100029.

(56) Zimm, B. H. Dynamics of Polymer Molecules in Dilute Solution: Viscoelasticity, Flow Birefringence and Dielectric Loss. The Journal of Chemical Physics 1956, 24, 269–278.

(57) Malevanets, A.; Kapral, R. Mesoscopic Model for Solvent Dynamics. The Journal of Chemical Physics 1999, 110, 8605–8613.

(58) Howard, M. P.; Nikoubashman, A.; Palmer, J. C. Modeling Hydrodynamic Interactions in Soft Materials with Multiparticle Collision Dynamics. Current Opinion in Chemical Engineering 2019, 23, 34–43.

(59) Gompper, G.; Ihle, T.; Kroll, D. M.; Winkler, R. G. In Advanced Computer Simulation Approaches for Soft Matter Sciences III ; Holm, C., Kremer, K., Eds.; Springer Berlin Heidelberg, 2009; pp 1–87.

(60) Allen, M. P.; Tildesley, D. J. Computer Simulation of Liquids; Oxford University Press, 1989.

(61) Ihle, T.; Kroll, D. M. Stochastic Rotation Dynamics: A Galilean-Invariant Mesoscopic Model for Fluid Flow. Phys. Rev. E 2001, 63, 020201.

(62) Nikoubashman, A.; Howard, M. P. Equilibrium Dynamics and Shear Rheology of Semi-flexible Polymers in Solution. Macromolecules 2017, 50, 8279–8289.

(63) Huang, C.; Chatterji, A.; Sutmann, G.; Gompper, G.; Winkler, R. Cell-level canonical sampling by velocity scaling for multiparticle collision dynamics simulations. Journal of Computational Physics 2010, 229, 168–177.

(64) Howard, M. P.; Panagiotopoulos, A. Z.; Nikoubashman, A. Efficient mesoscale hydrodynamics: Multiparticle collision dynamics with massively parallel GPU acceleration. Computer Physics Communications 2018, 230, 10–20.

(65) Ripoll, M.; Mussawisade, K.; Winkler, R. G.; Gompper, G. Low-Reynolds-number hydrodynamics of complex fluids by multi-particle-collision dynamics. Europhysics Letters (EPL) 2004, 68, 106–112.

(66) Mussawisade, K.; Ripoll, M.; Winkler, R. G.; Gompper, G. Dynamics of polymers in a particle-based mesoscopic solvent. The Journal of Chemical Physics 2005, 123, 144905.

