## Supporting Information for "Effect of Charge Distribution on the Dynamics of Polyampholytic Disordered Proteins"

| | SCD | nSCD |
|---|---|---|
| EKEKEKEKEKEKEKEKEKEKEKEKEKEKEKEKEKEKEKEKEKEKEKEKEK | -0.4131 | 0.000 |
| KEKKKEKEEEKKEKEKKEEKEKEEKKEEKKEEEKKKEEEKKEKKKEEEEK | -0.8686 | 0.017 |
| KEKKKEKEEKKEEKEKEEEKEEKEEKEKKKEEEKKKEKEEEKEKKKEKEK | -1.0960 | 0.025 |
| KEKKKEKEEEKEEEKKEEKEEEKEEKKKKKEEEKKKEKEEEKEKKKEKEK | -1.3237 | 0.033 |
| KEKEKEKKEEKEEEKEEEKEEEKKEKKEKKKEEEKKKEKKEEEKKKEKEK | -1.5513 | 0.041 |
| EEKKKKEEEKEEEEKKEEKEEEKKEKKEKKKEEEEKKEKKKEKKKEEKEK | -1.7793 | 0.050 |
| EEEKKKEEEKEEEKKKEEKEEEKKEEKEKKKKEEEKKKEKKEKKKEEKEK | -2.0067 | 0.058 |
| KKEEKEKEEEKEEEEEEKKKKEEKEKEKKKEKKEEEKKKEEKKKKEEEKK | -2.0347 | 0.059 |
| EKEEEEKEEEKKKKEEEKKEKEKEEKEKKKEKEEKEEKKKEKKKKKEEKE | -2.0698 | 0.060 |
| EEKKEEEEKEEEEKKKEEKKKEKKEKKKEKEEKKKKKEKEEEEEKKEKEK | -2.1291 | 0.063 |
| EEKKEEEEKEEEEKKKEEKKEEKKKKEKEKEEEKKKKEKKEEKEKKEKEK | -2.1581 | 0.064 |
| EKEKEEEEKEEEEKKKEEKKEEKKKKKEEKEKEKKKKKEKEEKEEKEKEK | -2.1866 | 0.065 |
| EKEKEEEEKEEEEKKKEEKKEEKKKEKEEKKKEKKKEKEKEKKEEKKKEE | -2.2170 | 0.066 |
| KKEEKEKKEEEEEKEEEKEEKKKEEEEKKKKKKKKEKKEKEEEKEKKEEK | -2.2417 | 0.067 |
| EEKEKEEKEKEEEEKKEKEEKKEEEEEKKKKKKKKEEKKKKKKEEKEKEE | -3.1560 | 0.100 |
| EEKEKEEKEKEEEEKEEKEEKKEEEEKKKKKKKKKEEKKKKKKEEKKEEE | -4.0707 | 0.133 |
| EEKEKEKEEKEEKEKEEEEEEKEEEEKKKKKKKEKEEKKKKKEKEKKEKK | -4.9860 | 0.167 |
| EEKKEKEEEEEEKEKEEEEEEKEEKEKKKKKKKKEKEKKKEKKKEKEEKK | -5.8993 | 0.200 |
| KEEEEKEEEKEEEEEEEKEKEKEEKEKKKKKKKKEKEKKKKKEKEKEKEK | -6.8134 | 0.233 |
| EEEEKEEKEEEEKEEEEEKKEKEEKEKEKKKKKKKEEKKKKKKEKKEKEK | -7.7272 | 0.267 |
| EEKEEEEKEEEEKEEEEEKKEKEEKEKEKKKEKKEEKKEKKKKKKEKKKK | -8.6394 | 0.300 |
| EEEKEKEKEKEEEEEEEEEEEKEEKEKEKKKKKKEEKKKEKKKKKEKKKK | -9.5552 | 0.333 |
| EKEKEKEEEEEEKEEEEEEEEKEEKEKEKKKKKKEEEKEKKKKKKKKKKK | -10.4666 | 0.367 |
| EKEKEEEEEEKEEEEEEEEKEEEKKEKEKKKKKKEEEKEKKKKKKKKKKK | -11.3868 | 0.400 |
| EKEEEEKEEEKEEEEEEEEKEEEKKEKEEKKKKKEEKEKKKKKKKKKKKK | -12.3037 | 0.434 |
| EEEEEEEEEEKEEEEEKEEKKKEKKEKEEEKKKKEEKEKKKKKKKKKKKK | -13.2435 | 0.468 |
| EEEEEEEEEEKEEEEEEEEKKKEKKKKEEEKKKKEKKEKKEKKKKKKKKK | -14.1189 | 0.499 |
| EEEEEEEEEEEEEEEEKEEKKKEKKKEKEEKKKKEKKEKKEKKKKKKKKK | -15.0046 | 0.532 |
| EEEEEEEEEEEEEEEEKEEKKEEKKKEEKKKEEKKKKEKKKKKKKKKKKK | -15.9407 | 0.566 |
| EEEEEEEEEEEEEEEEKEEKKEKEEKKKEKKKEKEKKKKEKKKKKKKKKK | -16.9036 | 0.601 |
| EEEEEEEEEEEEEEEEKEEKKEEEKKKEEKKKKKKKEKKKKKKKKKKKKE | -17.7742 | 0.633 |
| EEEEEEEEEEEEEEEEKEEEKEKEKEKEKKKKKKKKEKKKKKKKKEKKKK | -18.6958 | 0.667 |
| EEEEEEEEEEEEEEEEKEEEEKKEKEKEKKKKEKKKKKKEKKKKKKKKKK | -19.5876 | 0.699 |
| EEEEEEEEEEEEEEEKEEEEEKKKEEEEKKKKKKKKKKEKKKKKKKKKKK | -20.5398 | 0.734 |
| EEEEEEEEEEEKEEEEEEEEEEKEKEEKKKKKKKKKKKKEKKKKKKKKKK | -21.4576 | 0.767 |
| EEEEEEEEEKEEEEEEEEEEEEEEKEKKKKKKKKKKKKKKKKKKKKKKKE | -22.3636 | 0.800 |
| EEEEEEEEEEEEEEEEEKEEEEEEKEKKKKKKKKKKKKKKKKKKKKKKKE | -23.2591 | 0.833 |
| EEEEEEEEEEEEEEEEEEEEEKEEEKKKKKKKKKKKKKKKKKEKKKKKKK | -24.1881 | 0.867 |
| EEEEEEEEEEEEEEEEEEEEEKEKEKEKEKKKKKKKKKKKKKKKKKKKKK | -25.1497 | 0.902 |
| EEEEEEEEEEEEEEEEEEEEEEEKKKEKEKKKKKKKKKKKKKKKKKKKKK | -25.9326 | 0.930 |
| EEEEEEEEEEEEEEEEEEEEEEEEKKKEKKKKKKKKKKKKKKKKKKKKKK | -26.9742 | 0.968 |
| EEEEEEEEEEEEEEEEEEEEEEEEEKKKKKKKKKKKKKKKKKKKKKKKKK | -27.8421 | 1.000 |

Figure S1: Initially generated 42 E–K sequences for chain length $N = 50$ out of which 15 were chosen for detailed investigation of their conformational and dynamical properties. The chosen 15 E–K variants (EKVs) are shown on gray background. The last two columns show the SCD and the corresponding normalized SCD (nSCD) values for each of the 42 sequences.

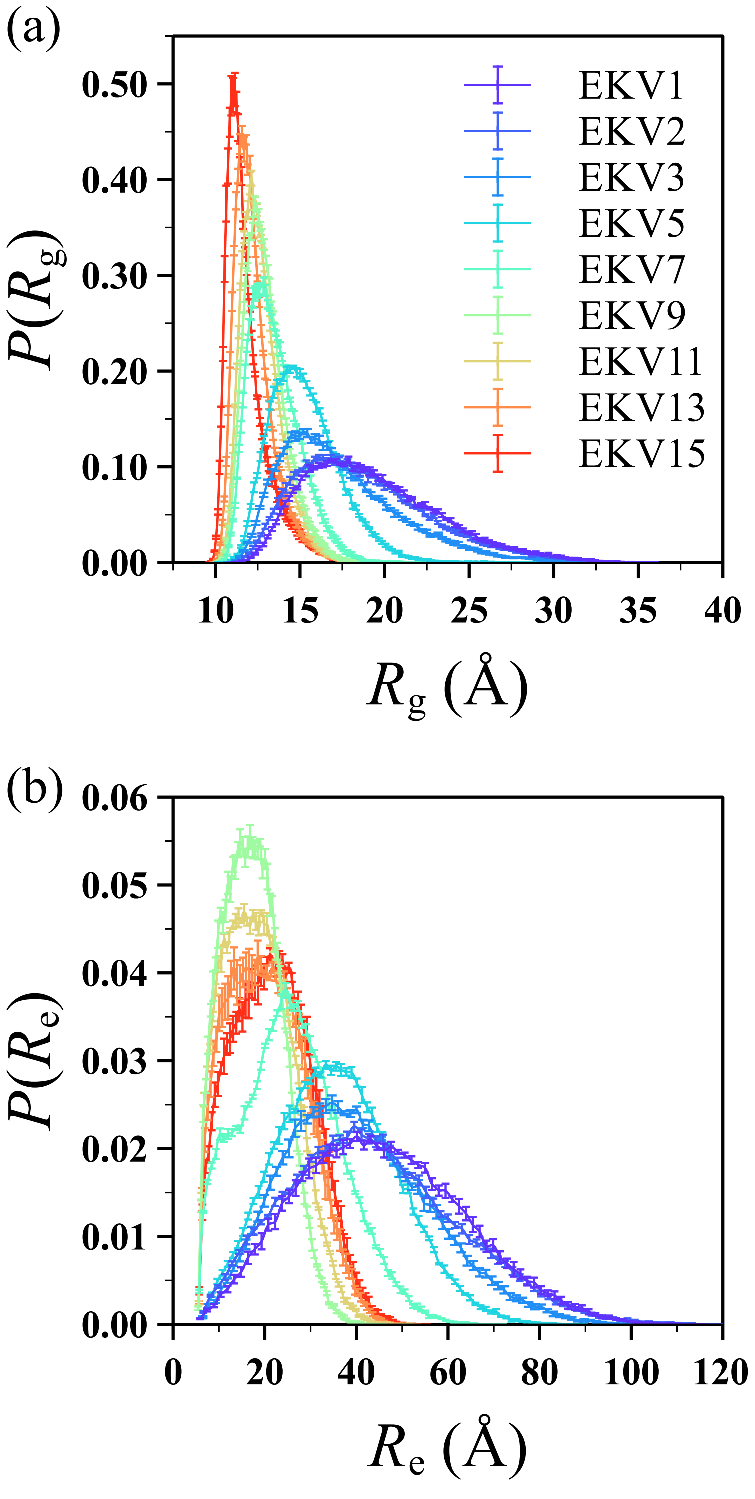


Figure S2: Probability distribution of (a) radius of gyration $R_{\mathrm{g}}$ and (b) end-to-end distance $R_{\mathrm{e}}$ for select EKVs.

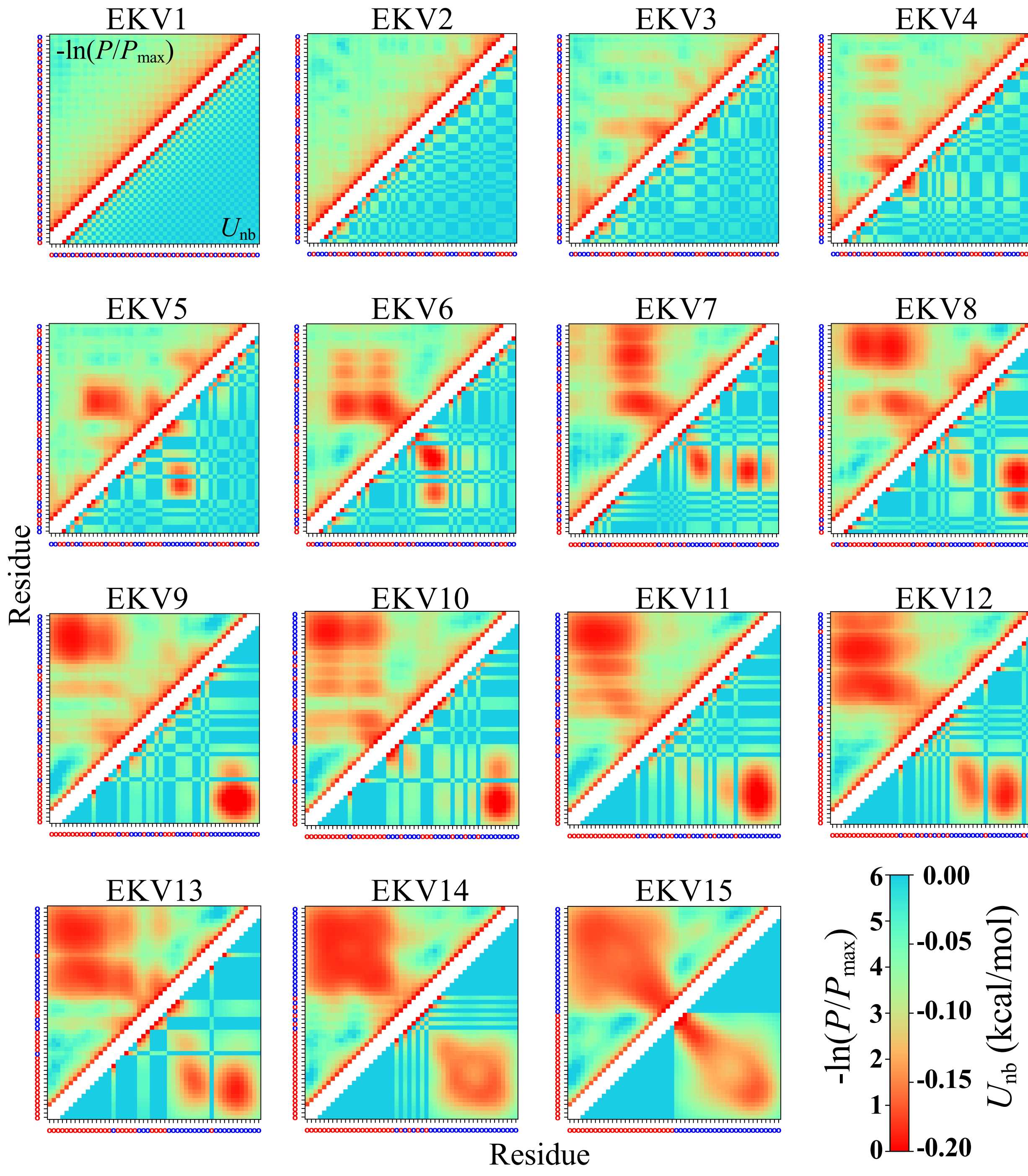


Figure S3: Interresidue distance-based contact map $-\ln(P/P_{\max})$ (above diagonal) and energy map $U_{\mathrm{nb}}$ (below diagonal) for all EKVs. The residue at each position is shown as a red (E) or blue (K) circle, respectively, on the axes. Two diagonals on either side of the main diagonal are removed in the maps to exclude directly bonded residues and residues that are separated by two bonds.

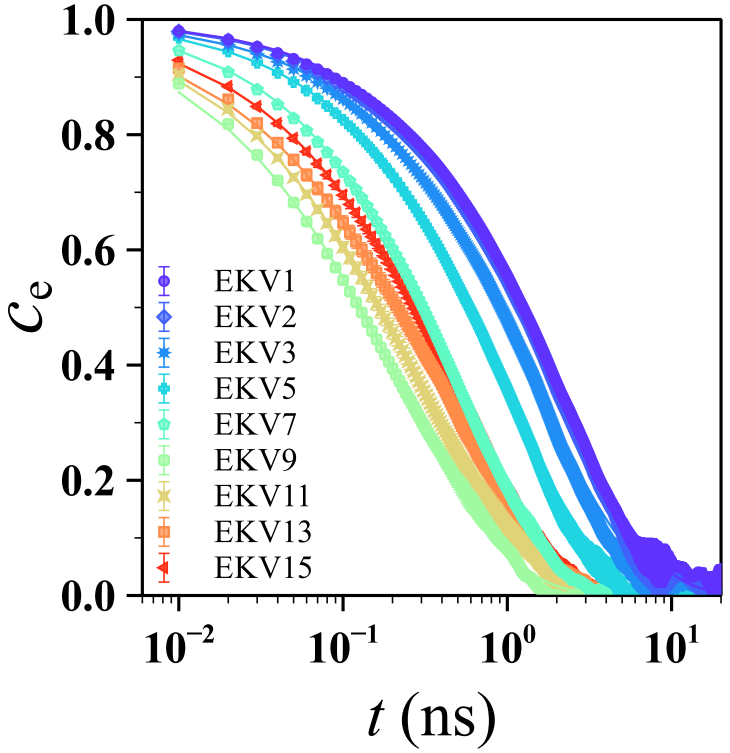


Figure S4: End-to-end vector autocorrelation function $c_e$ as a function of time $t$ for select EKVs.

| EKV | | SCD | nSCD |
|---|---|---|---|
| 1 | EKEKEKEKEKEKEKEKEKEKEKEKEKEKEK | -0.4218 | 0.000 |
| 2 | KEEEKKEEKKKEKEEKEEEKKEKEKEEKKK | -0.8964 | 0.038 |
| 3 | KEKKEEKEEEEEEKKKKEEKKKEKEKKEEK | -1.3694 | 0.075 |
| 4 | KKEEEKEEEEEEKKKKKKKKEKEEKKEEKE | -2.0027 | 0.125 |
| 5 | EKKEEEEEEEEEKKKKKKEKEKKEKKKKEE | -3.1086 | 0.212 |
| 6 | EEEKEEEEEEKEKEKKKKKKKEKEEKKKEK | -4.0586 | 0.288 |
| 7 | EEEKEEEEEEEEKKEKKKKKKKEKKEKEKK | -5.4191 | 0.395 |
| 8 | EEEEEEEKEEEEKKEKKEKKKKKKKEKKKE | -6.2656 | 0.462 |
| 9 | EEEEEEEEKEKEKEKEKEKEKEKKKKKKKK | -6.4684 | 0.478 |
| 10 | EEEEEEEEKEEEKKEKEKEKKKKKEKKKKK | -7.2072 | 0.537 |
| 11 | EEEEEEEEEEKEEKKEKKEEKKKKKKKKKK | -8.9930 | 0.678 |
| 12 | EEEEEEEEEEEKEKEKEEKKKKKKKKKKKK | -10.3674 | 0.787 |
| 13 | EEEEEEEEEEEEEKEKKEKKKKKKKKKKKK | -11.7251 | 0.894 |
| 14 | EEEEEEEEEEEEEEKKEKKKKKKKKKKKKK | -12.3565 | 0.944 |
| 15 | EEEEEEEEEEEEEEEKKKKKKKKKKKKKKK | -13.0670 | 1.000 |

Figure S5: Fifteen E–K variants (EKVs) for chain length $N = 30$ with their identifying number, SCD, and normalized SCD.

| EKV | | SCD | nSCD |
|---|---|---|---|
| 1 | KEKEKEKEKEKEKEKEKEKEKEKEKEKEKEKEKEKEKEKEKEKEKEKEKEKEKEKEKEKEKEKEKEKEKE | -0.4083 | 0.000 |
| 2 | EEKKKKEEEKKEEEKKEEEEKKKEEEKKEEKKEEEEKKEEKKKEKKEEKKEKKEKEKKKKEEEKEEEKKK | -1.5473 | 0.025 |
| 3 | KEKKEEEEEEKKEKKKKEEEKEKEEEEEKEKKEEEEKKEKKKKKKKEKKEEEKEKEKEKKKEEKEKEKEK | -2.1166 | 0.037 |
| 4 | EEKEKEKEKEEKEKEKEEKEEEEEEEEKEEKKKKKKKKKEKKEKEKEKKEKEKEKEKEKEKKEKEKEKEK | -3.5718 | 0.069 |
| 5 | EEKEKEKEKEEKEKEKEEKEEEEEEEEEEKKKKKKKKKKEKKEKEKEKKEKEKEKEKEKEKEKEKKEKEK | -3.8307 | 0.075 |
| 6 | KKEEEKEEKEEEEEEKKEEKEEEEEEEEEKKKKKKKKKKKEKEKKEEKKEEKKEKEKKKEKEEEKKKEKK | -5.8187 | 0.119 |
| 7 | EEEKEKEEKEKEKEEKEEEEEEEEEEEKEEKKKKKKEKKKKKKEKKEKKEKEKEEKKKEKEKEKKKEKEK | -7.2411 | 0.150 |
| 8 | EEEEEKEEKKEEKEEKKEEEEEEEEEEEEEKKKEKKEKKKKKKKKKKEKEKKEKKEKKEKEEKKKKEEKK | -9.8049 | 0.206 |
| 9 | EEEEKEKEEEEKEEEEEEEEEEEEEEKEEEKEKKEKKKKKEKKEKKKKKKKKKKKKEEKKEEEKKKKKKK | -18.6332 | 0.400 |
| 10 | EEEEEEEEEEEEEEEEEEKEKEKEKEKEKEKEKEKEKEKEKEKEKEKEKEKEKKKKKKKKKKKKKKKKKK | -22.1859 | 0.478 |
| 11 | EEEEEEEEEEKEEEEEEEEEEEEKEKEKKEEEEKKEKKKKEEKKKKKKKKKKKKKKEKEKKEKKKEKKKK | -22.9060 | 0.494 |
| 12 | EKEEEEEEEEEEEEEEEEEEEEEKEKEEKKEEEKKKEKKEKEKKKKEKKKKKKKKKKEKKKKKKEEKKKK | -23.7561 | 0.512 |
| 13 | EKEEEEEEEEEEEEEEEEEEEEEEEEKKEEKKEEKEKKEKKKEKEKKKKKKKKKKKKKKKEKKKEKKKKK | -28.0295 | 0.606 |
| 14 | EEEEEEEEEEEEEEEEEEEEEEEEEEEEEKEEEEKKEKKKEKKKKKKKKKKKKKKKKKKKKKKKKKKKKK | -42.9117 | 0.933 |
| 15 | EEEEEEEEEEEEEEEEEEEEEEEEEEEEEEEEEEEKKKKKKKKKKKKKKKKKKKKKKKKKKKKKKKKKKK | -45.9681 | 1.000 |

Figure S6: Fifteen E–K variants (EKVs) for chain length $N = 70$ with their identifying number, SCD, and normalized SCD.

| | SCD | nSCD |
|---|---|---|
| EEEEKEEEEEEEEEEEEEKEKEEEKKKKKKKKKKKKKKEKKKKEKKEEKK | -14.1025 | 0.499 |
| EEEEEKEEEEEEEEEEEEKEKEEEKKKKKKKKKKKKKKKEKEKKKEKEKK | -14.0387 | 0.497 |
| EEKEEEEEEEEEEEEEEEKKEEEEKKKKKKKKKKKKKKKEKEKEKKKEKK | -13.9012 | 0.492 |
| EEEEKEEEEEEEEEEEEEKKEEEEKKKKKKKKKKKKKKEKKEKKEKEKKK | -13.8417 | 0.490 |
| EEEEKEEEEEEEEEEEEKEKEEEEKKKKKKKKKKKKKKKEEKKKEKKKEK | -13.7130 | 0.485 |
| EEEEKEEEEEEEEEEEEKKEEEEEKKKKKKKKKKKKKEKKKEKKKEKEKK | -13.5460 | 0.479 |
| EEEEKEEEEEEEEEEEKEKEEEEEKKKKKKKKKKKKKKEKKEKKEKEKKK | -13.3372 | 0.471 |
| EEEEKEEEEEEEEEEEKKEEEEEEKKKKKKKKKKKKKKEKKKEKKEKEKK | -13.2434 | 0.468 |
| EEEEKEEEEEEEEEEEKKEEEEEEKKKKKKKKKKKKKKEKKEKKEKEKKK | -13.2135 | 0.467 |
| EEEEEKEEEEEEEEEKEKEEEEEEKKKKKKKKKKKKEKKKEKKKEKEKKK | -13.1051 | 0.462 |

Figure S7: E–K sequences for chain length $N = 50$ with their SCD and normalized SCD. Unlike EKV9 and EKV10 whose nSCD values are in the similar range (nSCD $\approx 0.5$), both the blocks of E and K residues do not appear at the termini in these sequences.

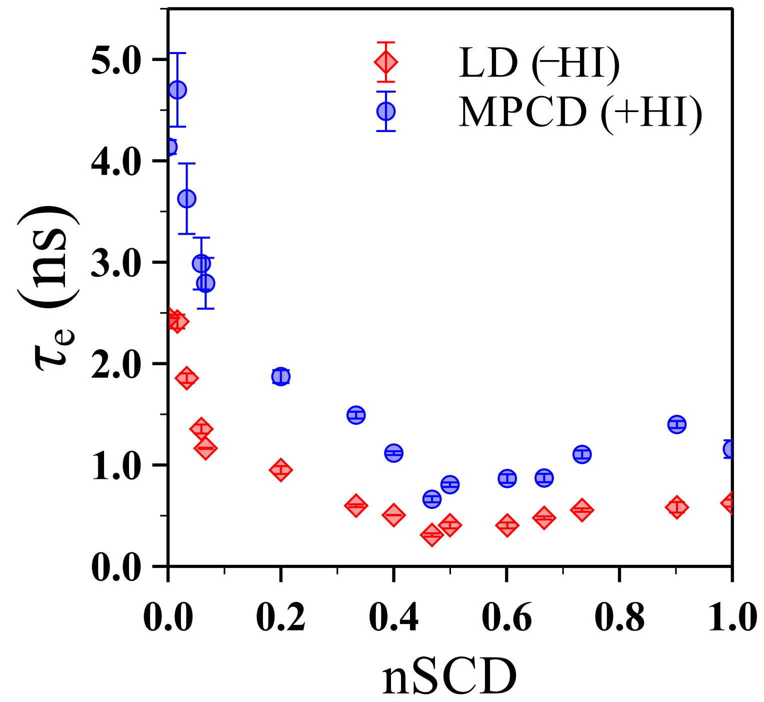


Figure S8: Relaxation time $\tau_e$ of the end-to-end vector obtained from the LD (−HI) and MPCD (+HI) simulations.
